# Atypical memory B cells form a pre-plasmacellular reservoir for steady-state IgD responses to common nasopharyngeal antigens

**DOI:** 10.1101/2023.08.29.554748

**Authors:** Roser Tachó-Piñot, Habib Bashour, Martyna Filipska, Sonia Tejedor-Vaquero, Leire de Campos-Mata, Alba Sáez-Gordón, Júlia Perera-Bel, Mauricio Guzman, Xavi Marcos-Fa, Pablo Canales-Herrerias, Jorge Domínguez-Barragán, Berta Arcós-Ribas, Andrei Slabodkin, Maria Chernigovskaya, María Luisa Rodríguez de la Concepción, José Gutierrez-Marcos, Ana García-García, Andrés Nascimento-Osorio, Mariona Pascal, Laia Alsina, Juan I. Aróstegui, Saurabh Mehandru, Charlotte Cunningham-Rundles, Jorge Carrillo, Giuliana Magri, Victor Greiff, Andrea Cerutti

**Affiliations:** Translational Clinical Research Program, Hospital del Mar Research Institute; Barcelona, Spain; School of Life Sciences, University of Warwick; Coventry CV4 7A, UK; Department of Immunology, University of Oslo and Oslo University Hospital; Oslo, Norway; Bioinformatics Unit, Research Programme on Biomedical Informatics, MARGenomics, Scientific & Technical Services, Institute “Hospital del Mar” for Medical Investigations (IMIM); Barcelona, Spain; Precision Immunology Institute, Icahn School of Medicine at Mount Sinai; New York, USA; Henry D. Janowitz Division of Gastroenterology, Department of Medicine, Icahn School of Medicine at Mount Sinai; New York, USA; IrsiCaixa AIDS Research Institute, Campus Can Ruti; Badalona, Spain; Pediatric Allergy and Clinical Immunology Dept., Clinical Immunology and Primary Immunodeficiencies Unit, Hospital Sant Joan de Déu; Barcelona, Spain; Study Group for Immune Dysfunction Diseases in Children, Institut de Recerca Sant Joan de Déu; Barcelona, Spain; Clinical Immunology Unit, Hospital Sant Joan de Déu-Hospital Clínic; Barcelona, Spain; Neuromuscular Pathology Unit, Neurology Service, Hospital Sant Joan de Déu-Hospital Clínic Barcelona; Spain; Immunology Service, Biomedical Diagnostics Center, Hospital Clinic; Barcelona, Spain; Institut d’Investigacions Biomèdiques August Pi i Sunyer (IDIBAPS); Barcelona, Spain; Departament of Medicine, Universitat de Barcelona; Barcelona, Spain; RETICS Asma, reacciones adversas y alérgicas (ARADYAL) and RICORS Red De Enfermedades Inflamatorias (REI), Instituto de Salud Carlos III; Madrid, Spain; Deptartment of Surgery and Surgical Specializations, Facultat de Medicina i Ciències de la Salut, University of Barcelona; Barcelona, Spain; Departments of Medicine and Pediatrics, The Prism Immunology Institute, The Icahn School of Medicine at Mount Sinai; New York, USA; Germans Trias i Pujol Research Institute (IGTP), Campus Can Ruti; Badalona, Spain; CIBERINFEC. Instituto de Salud Carlos III; Madrid, Spain; Immunology Unit, Department of Biomedical Sciences, Faculty of Medicine and Health Sciences, University of Barcelona; Barcelona, Spain; Department of Medicine, Icahn School of Medicine at Mount Sinai; New York, USA; Catalan Institute for Research and Advanced Studies (ICREA); Barcelona, Spain

## Abstract

The human nasopharyngeal mucosa includes organized lymphoepithelial structures continually engaged in frontline immune responses to aerodigestive antigens. Advancing our understanding of these responses might lead to the development of new strategies for the prevention and treatment of common immune disorders such as allergies. Here we identified a hitherto elusive tonsillar subset of atypical IgD class-switched IgD^+^IgM^-^ memory (IgD-ME) B cells that were clonally related to IgD^+^IgM^−^ germinal center (IgD-GC) B cells and IgD-secreting IgD^+^IgM^−^ plasma cells (IgD-PCs) but not anergic IgD^+^IgM^−^ B cells. Consistent with their pre-plasmacellular properties, IgD-ME B cells served as preferential precursors of IgD-PCs over IgD-GC B cells. IgD antibodies from IgD^+^IgM^−^ cells acquired reactivity to multiple oral, airborne and commensal antigens through a mutation-dependent pathway involving both innate and adaptive signals. Thus, IgD-ME B cells may form a ready-to-use pre-plasmacellular reservoir for steady-state IgD responses likely aimed at enhancing nasopharyngeal homeostasis.

**One Sentence Summary:** Tonsillar atypical memory B cells form a ready-to-use pre-plasmacellular repertoire for IgD responses to common aerodigestive antigens.

## INTRODUCTION

The human nasopharyngeal mucosa includes palatine, pharyngeal, lingual, and tubal tonsils, which are organized lymphoepithelial structures that constitute the human equivalent of the murine nasal-associated lymphoid tissue (*1*). Fissure-like openings on the tonsillar surface, termed crypts, convey commensal, airborne and food antigens from the lumen of the crypt to the underlying stratified epithelium and lamina propria, which are inhabited by abundant immune cells, including IgD-PCs (*1*). These PCs are thought to differentiate from IgD class-switched germinal center (IgD-GC) B cells undergoing IgM-to-IgD class switch recombination (CSR) and somatic hypermutation (SHM) in local lymphoid follicles (*2*).

In both humans and mice, CSR from IgM to IgD occurs via an unconventional pathway that initiates IgD responses to both autologous and microbial antigens (*3–7*). In humans but not mice, the IgD-GC B cells initiating these IgD responses express hypermutated Ig heavy (H) and light (L) chain genes (*2*) and differentiate to IgD-PCs with biased usage of IgL chains of the type lambda (Igλ) (*3*, *8*). Due to their pronounced autoreactivity and inability to further switch to downstream isotypes, IgD class-switched B cells have been proposed to have a tolerogenic origin, serving as a “sink” for autoreactive B cells (*5*, *9*).

While tonsillar IgD-GC B cells are thought to constitute the major precursors of IgD-PCs (*2*, *8*), direct evidence of the clonal relationship between IgD-GC B cells and IgD-PCs remains limited at best. It is also unclear whether tonsillar IgD-GC B cells generate IgD^+^IgM^−^ memory (IgD-ME) B cells and whether these IgD-ME B cells can further differentiate into IgD-PCs. In this regard, IgD-ME B cells have only been reported in the general circulation (*5*, *10*) and their phenotypic, transcriptional, and molecular landscapes, including clonal relationship with IgD-PCs, are as yet unknown. It is also unknown whether IgD responses involving tonsillar IgD-PCs target environmental antigens commonly present in the nasopharyngeal mucosa, including commensal antigens and potential allergens such as food and airborne proteins.

Here we found that IgD class-switched IgD^+^IgM^−^ B cells from human tonsils included IgD-ME B cells with atypical phenotypic and transcriptional signatures, unique molecular architecture, extensive clonal affiliation with both IgD-GC B cells and IgD-PCs, and pronounced pre-plasmacellular properties. Accordingly, IgD-ME B cells clonally differentiated into IgD-PCs via a complex mutation-intensive pathway that reflected sustained antigenic stimulation of both IgD-ME B cells and their IgD-GC precursors over time. This pathway involved multiple innate and adaptive signals and induced IgD responses to multiple oral, airborne and commensal antigens through a mutation-dependent mechanism that enhanced IgD secretion in response to inflammatory signals, including T helper type-2 (T_H_2) cell-driven signals.

## RESULTS

### IgD-PCs inhabit follicular in addition to epithelial compartments from human tonsils

To determine whether tonsillar IgD-GC B cells generate IgD-ME B cells capable of further differentiating into IgD-PCs, we first analyzed human tonsils by multi-parametric tissue immunofluorescence analysis (IFA). IgD-PCs with large cytoplasm, eccentric nucleus, and abundant intracellular IgD but no intracellular IgM inhabited both crypt epithelium and sub-epithelial areas (**Fig. 1A**), which are enriched in memory B cells (*11*). Of note, some tonsillar GCs included an abundant and morphologically heterogeneous population of IgD class-switched IgD^+^IgM^−^ cells that likely encompassed PCs in addition to both GC and post-GC B cells (**Fig. 1B**). By showing that IgD-PCs inhabit both lymphoid inductive and epithelial effector sites of tonsils, these observations suggest that tonsillar IgD-GC B cells may generate IgD-ME B cells in addition to IgD-PCs.

**Fig. 1.**
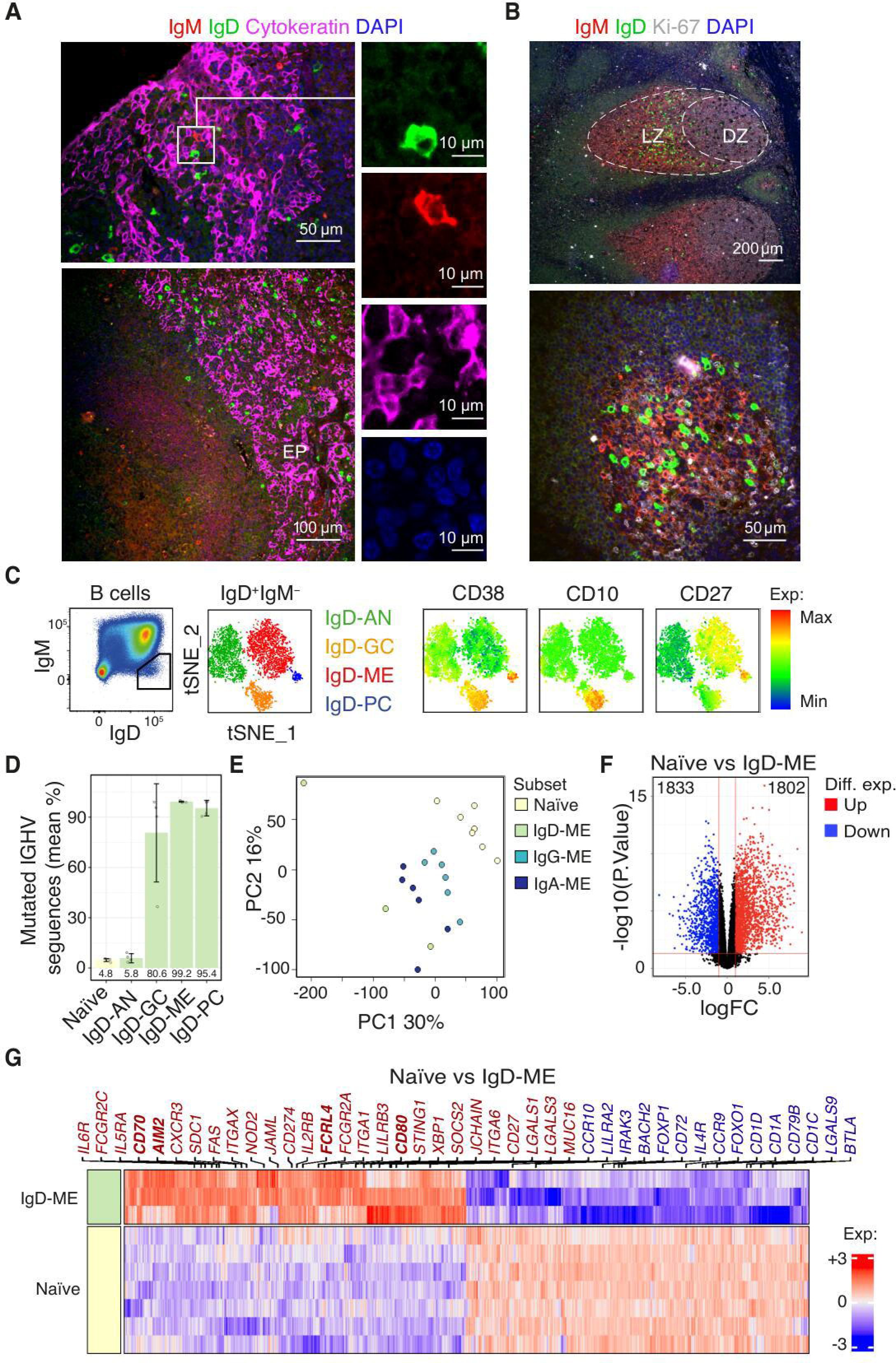
Tonsillar IgD class-switched B cells inhabit both lymphoid and epithelial compartments and include IgD-ME B cells in addition to IgD-GC B cells and IgD-PCs. **(A)** Immunofluorescence analysis of human tonsillar crypt epithelium stained for IgD (green), IgM (red), cytokeratin (magenta), and nuclear DNA (blue). Inset (top left panel) includes intraepithelial IgD-PCs and control IgM-PCs, which are further visualized after digital magnification and no color merging (right panels). (B) Immunofluorescence analysis of human tonsillar lymphoid follicles stained for IgD (green), IgM (red), Ki-67 (white), and nuclear DNA (blue). Dashed line (top panel) demarcates the GC of a lymphoid follicle. LZ, light zone; DZ, dark zone; M, mantle. (C) Flow cytometry of total tonsillar IgD^+^IgM^−^ B cells with tSNE plot and clusters defined based on the expression of CD10, CD38, CD27, Igλ, CD11c, CD43, CD21, CD95, CXCR3 and CD69. Plots on the right reflect relative CD38, CD10 and CD27 expression. Representative data from three independent experiments. (D) Mean proportion of IGHV mutated sequences across IgD-ME B cells, IgD-GC B cells and IgD-PCs as well as control naïve B cells and anergic IgD-AN B cells from human tonsils (n = 4 per cell type). Numbers below bars represent the mean percentage across donors. Error bars represent SD. (E) Principal component analysis (PCA) of RNA-sequencing transcriptome from Naïve, IgD-ME, IgG-ME and IgA-ME B cells sorted from human tonsils. (F) Volcano plot summarizing gene fold change (logFC) and adjusted P value, as −log10(padj), between Naïve and IgD-ME B cells as in (E). (G) Heatmap summarizing top 1418 differentially expressed genes between naïve (n=7) and IgD-ME B cells (n=3) as in (E). Highlighted genes of interest are upregulated (red) or downregulated (blue) in IgD-ME B cells compared to naïve B cells. The color bar depicts normalized intensity values. Genes highlighted in bold are discussed in the text.

### Tonsillar IgD-ME B cells coexist and share GC molecular traits with IgD-PCs

IgD-ME B cells have been described in the peripheral circulation (*5*, *10*), but their existence in tonsils and other lymphoid tissues remains elusive. To identify tonsillar IgD-ME B cells, we first performed unbiased clustering of the global population of tonsillar IgD class-switched IgD^+^IgM^−^ B cells upon its flow cytometric characterization. Cluster visualization onto a T-distributed stochastic neighbor embedding (t-SNE) projection revealed the presence of four major subsets of IgD^+^IgM^−^ B cells, which were defined as CD10^−^CD27^hi^CD38^hi^ IgD-PCs, CD10^+^CD38^+^ IgD-GC B cells, CD10^−^CD27^−^CD38^low^ IgD anergic (IgD-AN) B cells, and putative CD10^−^CD27^+^CD38^−^ IgD-ME B cells (**Fig. 1C, Fig. S1A**). Of note, IgD-AN B cells have been previously described as non-class-switched, non-mutated and autoreactive naïve B cells that down-regulate IgM and are poorly responsive to external stimuli (*12*).

Unlike these IgD-AN B cells, IgD-ME B cells should have a GC-driven molecular configuration similar to that of IgD-GC B cells and IgD-PCs, which are highly mutated (*8*, *13*). To formally address this point, we adopted a standard flow cytometry strategy (**Fig. S1B**) and used it to sort-purify individual IgD^+^IgM^−^ cell clusters along with control naïve and IgD-AN B cells. High-throughput next-generation sequencing (NGS) of immunoglobulin heavy chain variable (IGHV) genes revealed that, similar to IgD-PCs and IgD-GC B cells, the majority of IgD-ME B cells were mutated, whereas IgD-AN B cells and naïve B cells were not (**Fig. 1D**).

To further validate the ME B cell nature of IgD-ME B cells, we analyzed the transcriptome of sort-purified tonsillar IgD-ME B cells and compared it to the transcriptome of naïve B cells as well as canonical IgG-ME and IgA-ME B cells. Principal component analysis (PCA) showed that IgD-ME B cells clustered together with IgG-ME and IgA-ME B cells but were clearly distinct from naïve B cells (**Fig. 1E**). Volcano plot and heatmap analyses showed that over three thousand genes were differentially expressed by IgD-ME and naïve B cells (**Fig. 1F**), *CD70, CD80*, *FCRL4*, *AIM2* and *MUC16* being hallmarks of ME B cells, including respiratory memory B cells (*14–17*), up-regulated by the IgD-ME subset (**Fig. 1G**). Thus, the human nasopharyngeal mucosa encompasses GC-derived IgD-ME B cells among other IgD^+^IgM^−^ B cell subsets.

### Tonsillar IgD-ME B cells exhibit atypical phenotypic, molecular, and functional properties

To elucidate the properties of IgD-ME B cells, we first defined them as IgD^+^IgM^−^CD10^−^CD19^+^CD38^−^CD27^+^ cells and then calculated their frequency within total ME switched IgM^−^CD10^−^CD19^+^CD38^+^ cells from different tissues. IgD-ME B cells were enriched in tonsils compared to splenic tissue (**Fig. 2A**), possibly indicating their preferential induction in the aerodigestive mucosa. IgD-ME B cells were also detected in the circulation (**Fig. 2A**), albeit at a somewhat lower frequency than in tonsils, which might reflect the transit of IgD-ME B cells from tonsillar inductive sites to systemic effector areas (*3*). Compared to naïve B cells, IgD-ME B cells showed TACI up-regulation and elevated Igλ usage (**Fig. S2A**), two hallmarks of IgD class-switched B cells (*3*).

**Fig. 2.**
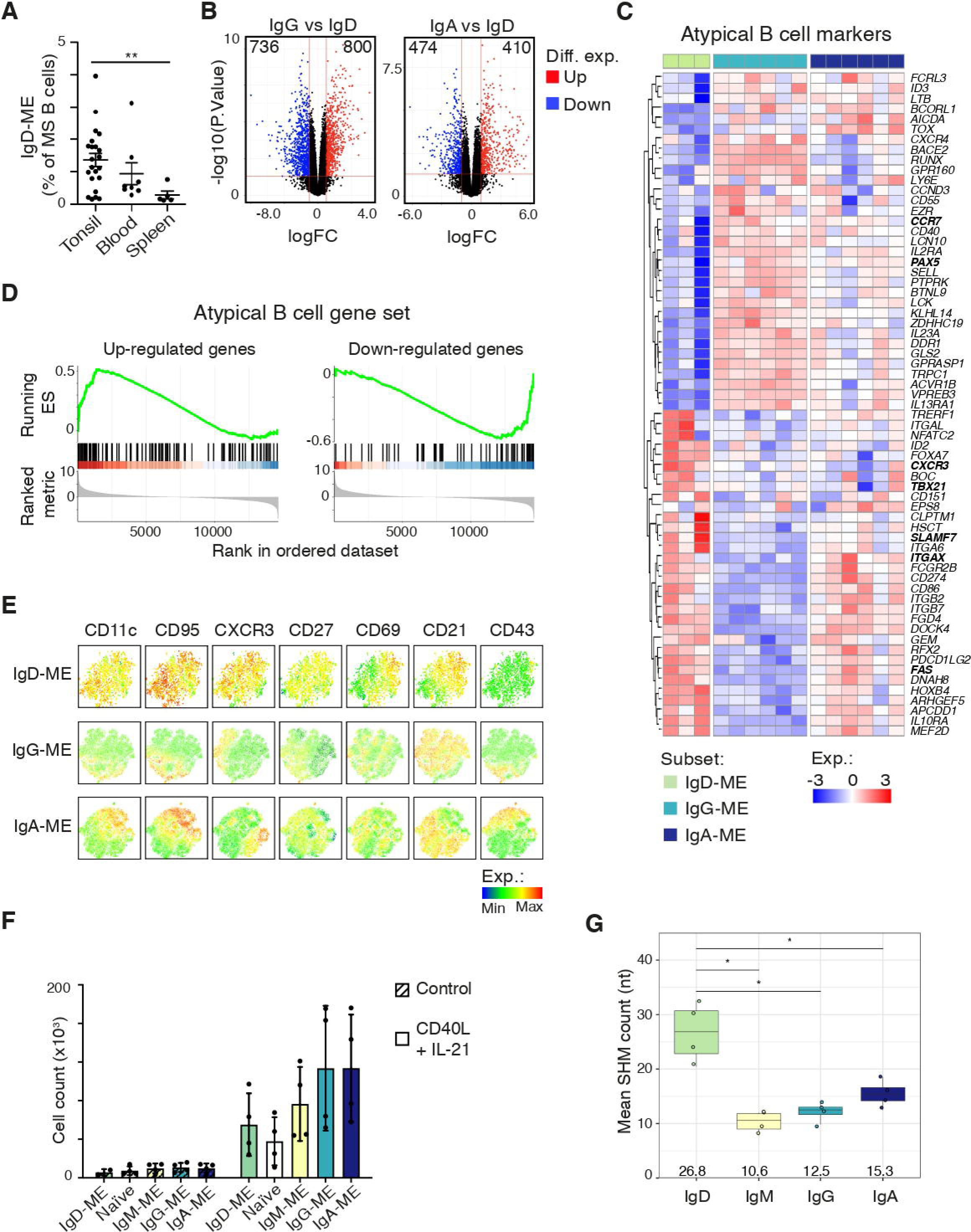
Tonsillar IgD-ME B cells exhibit phenotypic, mutational and transcriptional properties consistent with persistent exposure to and activation by mucosal antigens. (**A**) Frequency of IgD-ME B cells within total class-switched CD45^+^CD19^+^CD38^−^CD10^−^IgM^−^ ME B cells from human tonsils (n = 22), peripheral blood (n = 8) or spleen (n = 5). Error bars represent SD. (**B**) Volcano plot summarizing gene fold change (logFC) and adjusted P value, as - log10(padj), between IgD-ME and IgG-ME (left) or IgA-ME (right) B cells determined by RNA-sequencing of B cell subsets sorted from human tonsils. Numbers in plots indicate differentially expressed genes (|logFC| > 1 and p.value < 0.05) up-or down-regulated on IgD-ME. (**C**) Heatmap depicting differentially expressed genes (adj.P.Value < 0.05 and |log2FC| > 1) between IgD-ME (n=3) vs. IgG-ME + IgA-ME (n=6 each) B cells belonging to a previously-described list of atypical B cell markers (Holla et al. 2021). The color bar depicts normalized intensity values. Genes highlighted in bold are discussed in the text. (**D**) Gene set enrichment analysis (GSEA) exploring enrichment of the atypical B cell signature described in (*18*), in IgD-ME B cells, when compared to IgG-ME + IgA-ME B cells. ES: enrichment score. (**E**) Expression intensity of CD11c, CD95, CXCR3, CD27, CD69, CD21 and CD43 molecules projected on tSNE plots of IgD-ME, IgG-ME or IgA-ME B cells from human tonsils, determined by spectral flow cytometry. Representative data from three independent experiments. (**F**) Absolute number of viable cells/well after culturing FACSorted Naïve, IgM-ME, IgD-ME, IgG-ME or IgA-ME B cells for 6 days with medium alone (control) or CD40L and IL-21. Data summarizes four independent experiments. (**G**) Mean somatic hypermutation (SHM) count across IgD/IgM/IgG/IgA-ME B cells from human tonsils (n=4). Numbers below boxes represent median values of the four data points. Differences were assessed with Kruskal-Wallis test followed by a post-hoc pairwise Dunn’s (A) or a Mann-Whitney (F-G) test. * p < 0.05, ** p < 0.01. Statistical significance (F-G) was only calculated between IgD-ME B cells and other subsets. Comparisons lacking statistical reporting are not statistically significant.

Volcano plots obtained from transcriptome analysis showed hundreds of differentially expressed genes (DEGs) in IgD-ME B cells compared to canonical IgG-ME or IgA-ME B cells (**Fig. 2B**), which pointed to distinct differentiation, signaling, homing and/or functional properties. DEGs included *LGALS1* (galectin-1), *FCGR2B* (FcR2B), and *CD274* (PD-L1), which were up-regulated in IgD-ME B cells and mediate immune regulation, as well as *CD79b*, *CD1*, and *CCR9*, which were down-regulated in IgD-ME B cells and mediate B cell activation, lipid presentation, and gut migration, respectively (**Fig. S2B**).

Compared to IgG-ME and IgA-ME B cells, tonsillar IgD-ME B cells exhibited a prominent transcriptional signature overlapping with that of atypical ME B cells (*18*), including up-regulation of *ITGAX* (CD11c), *CXCR3*, *SLAMF7*, *TBX21* (T-bet), and *FAS* (CD95). In addition, IgD-ME B cells showed down-regulation of *PAX5* and *CCR7* (**Fig. 2C**). Most of these DEGs showed similar expression patterns on IgD-ME B cells when compared to canonical IgG-ME and IgA-ME B cells by gene set enrichment analysis (GSEA) (**Fig. 2D**), suggesting IgD-ME B cells are enriched in an atypical memory B cell signature. Aside from confirming the up-regulation of CD11c, CD95, CXCR3, CD69, and FCRL4, flow cytometry showed the down-regulation of CD21 on tonsillar IgD-ME B cells (**Fig. S2C**), which is yet another hallmark of atypical memory B cells (*18*). Circulating and splenic IgD-ME B cells partly recapitulated this atypical phenotype (**Fig. S2D** and **S2E**).

Besides further corroborating these data, t-SNE projections highlighted discrete co-expression patterns within tonsillar ME B cell sub-populations. While the activation molecules CD11c and CD95 were mostly co-expressed by discrete subsets of IgD-ME and IgA-ME B cells and by a smaller but distinct fraction of IgG-ME B cells, the activation molecule CD43 was mostly expressed by overlapping fractions of IgG-ME and IgA-ME but not IgD-ME B cells (**Fig. 2E**). Of note, CD21 followed an opposite expression pattern compared to CD11c and CD95 on IgG-ME and IgA-ME but not IgD-ME B cells. Thus, IgD-ME B cells express transcriptional and phenotypic properties of atypical ME B cells, which are characterized by functional hyporesponsiveness to activation (*18*).

To test whether also IgD-ME B cells were functionally hyporesponsive, we compared their *in vitro* proliferation with that of control naïve B cells or canonical IgG-ME and IgA-ME B cells in a 6-day culture supplemented with CD40L and IL-21, which mimic B cell stimulation by antigen-activated T cells. IgD-ME B cells were found to proliferate slightly less than canonical IgG-ME or IgA-ME B cells but comparably to naïve or IgM-ME B cells (**Fig. 2F**).

In light of these findings, we hypothesized that tonsillar IgD-ME B cells could become overactivated as a result of their continuous exposure to antigen from the lumen of crypts, which could lead to extensive SHM due to iterative IgD-ME B cell entry into the GC reaction. Consistent with their possible GC-driven selection by antigen (*19*, *20*), IgD-ME B cells displayed a significantly higher mean mutation count compared to canonical ME B cell subsets (**Fig. 2G**) and more mutations in antigen-binding complementarity-determining regions (CDRs) (**Fig. S3A**), which also featured a higher replacement versus silent mutation ratio (R:S ratio) compared to framework regions (FWRs) (**Fig. S3B**). Thus, IgD-ME B cells constitute a unique tissue-based subset of possibly antigen-selected ME B cells with phenotypic and transcriptional properties of atypical ME B cells.

### Tonsillar IgD-ME B cells share a unique molecular architecture with IgD-PCs

We then compared the Ig V_H_DJ_H_ gene usage of IgD class-switched B cell subsets with that of GC B cells, ME B cells and PCs expressing IgM, IgG or IgA. Additional comparisons were made with naïve B cells co-expressing surface IgD and IgM and IgD-AN expressing surface IgD but not IgM. Similar to naïve B cells, IgD-AN B cells co-expressed IgD and IgM transcripts, in agreement with previous studies showing absence of IgD class-switching in this subset of B cells (*12*). This co-expression and the possibility that rare CD27^−^ B cells from the IgD-ME subset could contaminate naïve and IgD-AN B cells prompted us to analyze IgM rather than IgD transcripts from naïve and IgD-AN B cell samples, the latter being defined hereafter as IgM-AN. We found that IgD-GC B cells, IgD-ME B cells and IgD-PCs had a unique pattern of IGHV gene usage, which was distinct from that of other tonsillar B cell subsets, including B cells expressing IgG or IgA (**Fig. 3A**). Consistent with this, IgD class-switched cell subsets from all four donors clustered together and separate from other B cells (**Fig. 3A**).

**Fig. 3.**
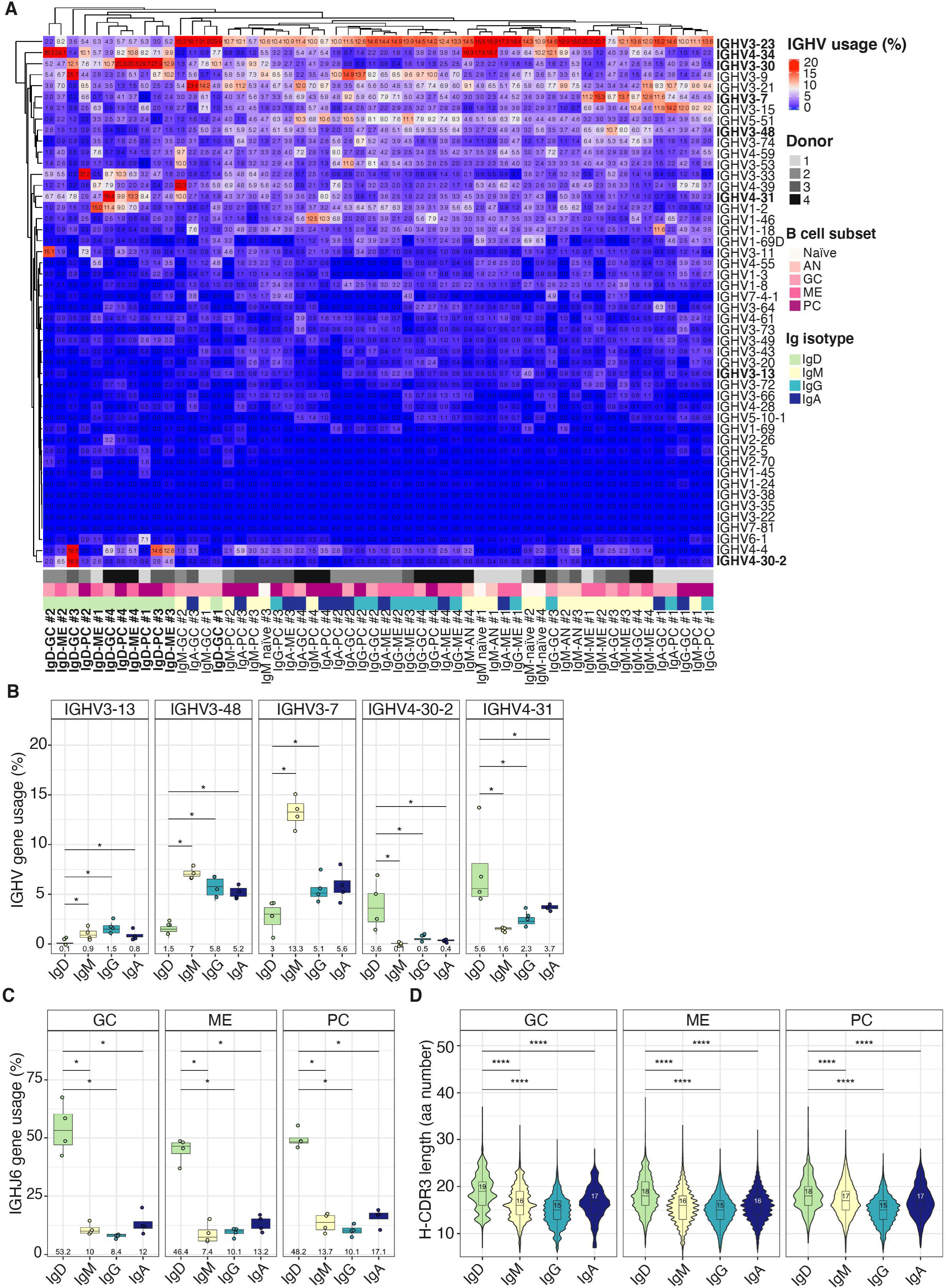
Tonsillar IgD-ME B cells express a unique IgD gene repertoire and a longer H-CDR3 segment similar to IgD-GC B cells and IgD-PCs. (**A**) V_H_ gene usage pattern across samples is shown as a heatmap where rows and columns represent IGHV genes and samples, respectively. Samples are color-coded by donor (1-4), cell subset (naïve, anergic, GC, ME or PC) and antibody isotype (IgM, IgD, IgG or IgA). Genes highlighted in bold are mentioned in the text. (**B**) Mean IGHV3-13, IGHV3-48, IGHV3-7, IGHV4-30-2 and IGHV4-31 gene usage by ME B cells from human tonsils expressing surface IgM, IgD, IgG or IgA alone (n = 4). (**C**) Mean IGHJ6 gene usage by ME B cells, GC B cells or PCs from human tonsils expressing surface IgD, IgM, IgG or IgA alone (n = 4). In (**B**) and (**C**), numbers below boxes represent median values of average gene usage. (**D**) Comparison of clonal CDR3H length (aa – amino acids) among antibody isotypes (IgM, IgD, IgG or IgA) expressed by B cell subsets (ME, GC or PCs) from human tonsils. Numbers in plots represent median CDR3 length. Differences were assessed with Kruskal-Wallis test followed by a post-hoc pairwise Mann-Whitney test with p-value adjustment following Benjamini-Hochberg method. * p < 0.05, **** p < 0.0001. Comparisons lacking statistical reporting are not statistically significant.

Compared to other ME B cells, IgD-ME B cells used less IGHV3-7, IGHV3-13 and IGHV3-48 but more IGHV4-30-2 and IGHV4-31 (**Fig. 3B**). IgD-GC B cells, IgD-ME B cells and IgD-PCs also showed a trend towards lower IGHV3-23 and higher IGHV3-30 and IGHV4-34 usage (**Fig. S3C**), as also noted in previously published but smaller scale studies (*9*, *21*). A subset (5-25%) of IgD-GC B cells, IgD-ME B cells and IgD-PCs used IGHV3-30 and IGHV4-34 in our dataset (**Fig. S3C**). In addition, IgD-GC B cells, IgD-ME B cells and IgD-PCs used more IGHJ6 than their IgM, IgG and IgA counterparts (**Fig. 3C**). Considering that GC B cells usually counter-select IGHV4-34 as well as IGHJ6 due to their intrinsic autoreactivity (*9*, *22*), these results suggest that tonsillar IgD responses may target a unique set of foreign epitopes structurally similar to self-antigens. Besides increased usage of IGHV4-34 and IGHJ6 genes, autoreactive and polyreactive antibodies have been previously associated with longer than average H-CDR3 regions (*9*, *12*). Accordingly, the length of H-CDR3 in IgD-GC B cells, IgD-ME B cells and IgD-PCs was higher than in other B cell subsets (**Fig. 3D**). Thus, tonsillar IgD-ME B cells emerge from a GC-driven response that further induces IgD-PCs and involves the selection of intrinsically autoreactive and polyreactive IGHV and IGHJ genes.

### Tonsillar IgD-ME B cells are clonally affiliated to IgD-PCs

Next, we ascertained the degree of clonal overlap amongst various IgD class-switched IgD^+^IgM^−^ subsets. A clone was defined as a cluster of B cell receptor (BCR) sequences utilizing a unique combination of IGHV and IGHJ genes and an identical CDR3 amino acid sequence. IgD-ME B cells clonally overlapped with IgD-GC B cells and IgD-PCs (**Fig. 4A**). This overlap was higher than the clonal overlap observed amongst other tonsillar B cell subsets. In addition, IgD-ME B cells as well as IgD-GC B cells and IgD-PCs only rarely clonally overlapped with equivalent tonsillar B cells or PCs expressing IgM, IgG or IgA, or with naïve or IgM-AN B cells (**Fig. 4A**).

**Fig. 4.**
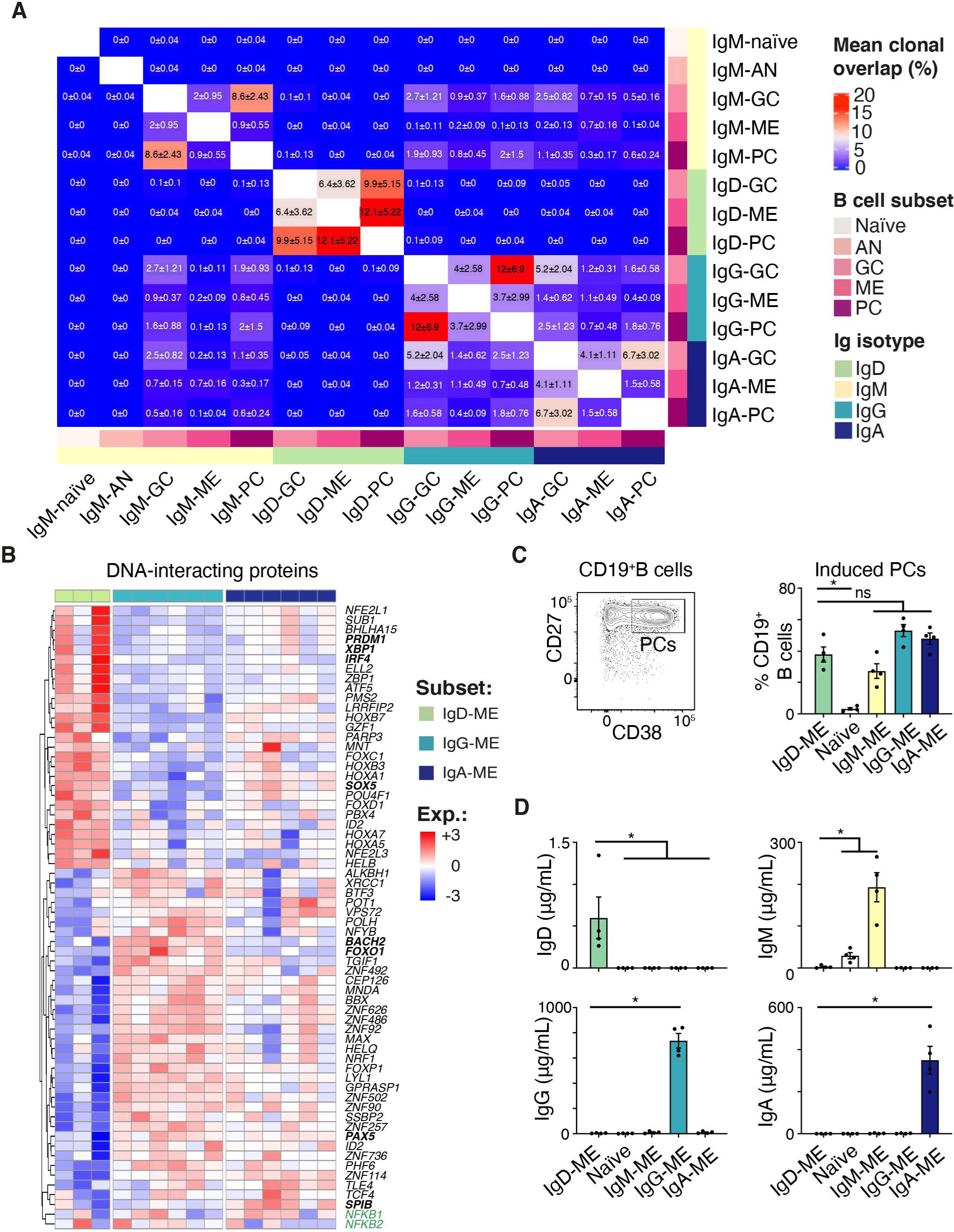
Tonsillar IgD-ME B cells are clonally affiliated to IgD-PCs and transcriptionally and functionally poised to become IgD-PCs. (**A**) Values of mean pairwise clonal overlap (%) across tonsillar B cell subsets (naïve, anergic, ME or GC B cells and PCs). Clones were considered overlapping when they had an identical CDR3 amino acid sequence and utilized the same V and J germline genes. Each cell within the heatmap contains the mean clonal overlap value ± SD (n = 4). (**B**) Heatmap showing relative expression of DEGs determined by RNA-seq (adj.P.Value < 0.05 and |log2FC| > 1) and encoding DNA-interacting proteins in tonsillar IgD-ME, IgG-ME, and IgA-ME B cells. The color bar depicts normalized intensity values. Genes highlighted in bold are discussed in the text; genes highlighted in green are mutated in primary immunodeficiencies and HIDS from Fig. 6 F and G. (**C**) Representative flow cytometry gating strategy and percentage of CD19^+^CD27^high^CD38^high^ PCs obtained after culturing FACSorted naïve or CD27^+^ memory B cells expressing surface IgM, IgD, IgG or IgA alone for 6 days with CD40L and IL-21. Data summarize four independent experiments. Error bars represent SD (**D**) ELISA of IgD, IgM, IgG or IgA secreted by naïve B cells or memory B cells cultured as in (B) (n=4). Differences were assessed with Kruskal-Wallis test followed by a post-hoc Mann-Whitney test. * p < 0.05. Comparisons lacking statistical reporting are not statistically significant. Error bars represent SD.

Having shown that IgD-ME B cells were clonally related to IgD-PCs, we considered the possibility that IgD-ME B cells may be prone to differentiate into IgD-PCs. When compared to IgG-ME and IgA-ME B cells, IgD-ME B cells showed several DEGs encoding DNA-interacting proteins relevant to PC differentiation, including up-regulated *PRDM1*, *XBP1*, and *IRF4*, and down-regulated *FOXP1*, *PAX5*, *FOXO1 BACH2*, and *SPIB* (**Fig. 4B**) (*23*, *24*). Accordingly, IgD-ME B cells induced as many PCs as other memory B cells upon 6-day culture with CD40L and IL-21 (**Fig. 4C**). Of note, only IgD-ME B cells yielded detectable IgD secretion and did not release IgG or IgA (**Fig. 4D**), which supported their role as IgD-PC intermediates as well as their lack of clonal overlap with IgG-ME or IgA-ME B cells. Thus, IgD-ME B cells uniquely clonally overlap with IgD-GC B cells and IgD-PCs, show an enriched PC transcriptional signature, and selectively differentiate to IgD-PCs.

### Tonsillar IgD-ME B cells exhibit pre-plasmacellular properties

We further examined the PC differentiation potential of IgD-ME B cells by interrogating their transcriptome for any enrichment in gene programs related to antibody secretion. When compared to canonical IgG-ME or IgA-ME B cells, IgD-ME B cells were enriched in gene sets linked to protein synthesis, post-translational modification, and trafficking processes involving the endoplasmic reticulum (ER) and Golgi apparatus (**Fig. 5A**). In particular, up-regulated genes included those involved in the unfolded protein response (UPR), a molecular program required for antibody secretion (*24*). Aside from being enriched in *ERN1* and *XBP1* (**Fig. 5B**), which are key to the UPR (*24*, *25*), IgD-ME B cells showed up-regulated *IL6R* and *IFNAR2* (**Fig. 5C**), which encode receptors for IL-6 and IFN-α cytokines involved in PC differentiation (*26*).

**Fig. 5.**
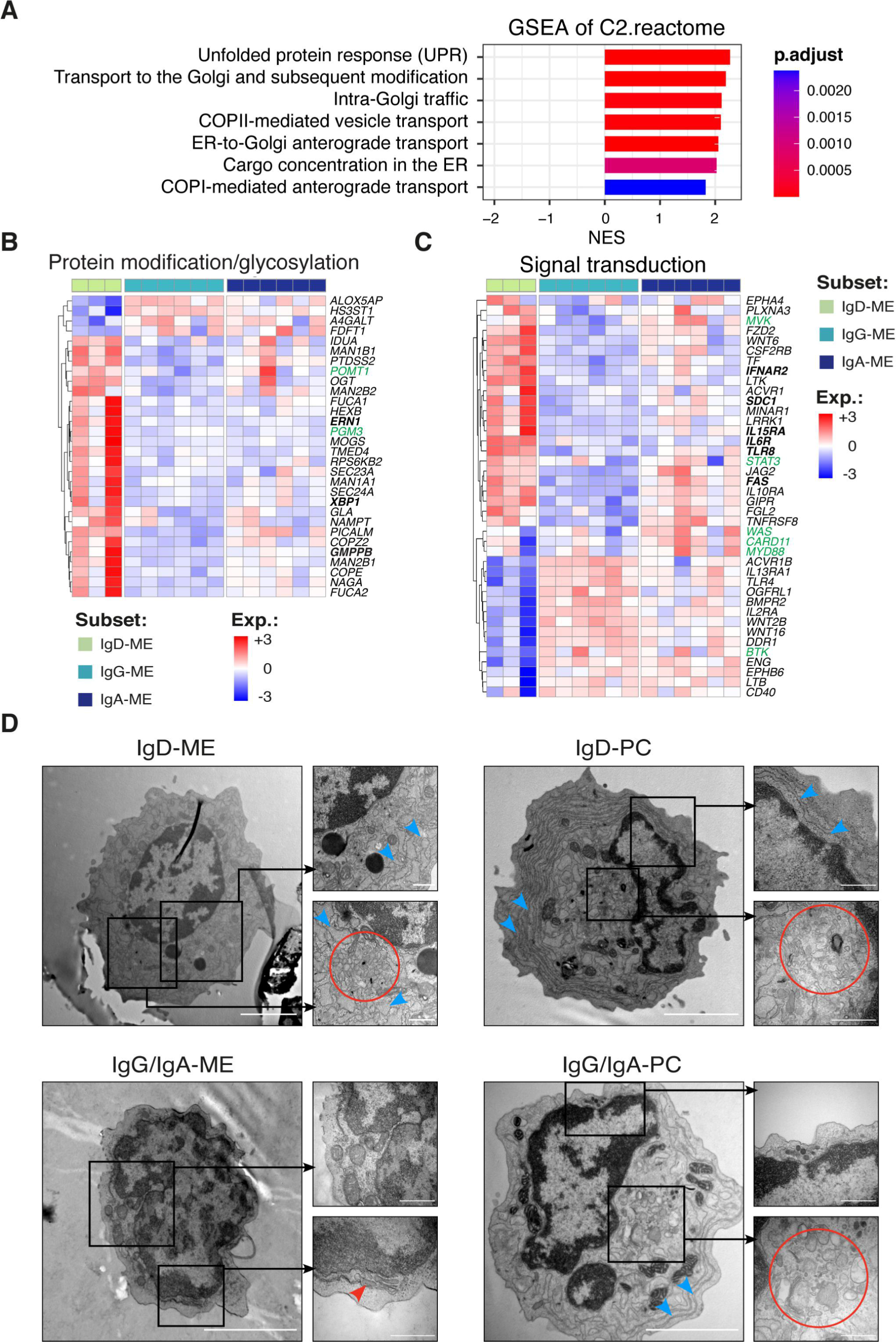
Tonsillar IgD-ME B cells show pronounced pre-plasmacellular properties. (**A**) Bar plot showing significantly enriched terms (p adj < 0.05) in gene set enrichment analysis (GSEA) from RNA-seq analysis of tonsillar IgD-ME B cells compared to IgG-ME plus IgA-ME B cells. NES; normalized enrichment score. (**B** and **C**) Heatmaps showing genes encoding protein modification/glycosylation factors (B) and signal transduction proteins (C) differentially expressed (adj.P.Value < 0.05 and |log2FC| > 1) by tonsillar IgD-ME vs. IgG-ME or IgA-ME B cells. The color bar depicts normalized intensity values. Genes highlighted in bold are discussed in the text; genes highlighted in green are mutated in primary immunodeficiencies or HIDS from Fig. 6F. (**D**) Transmission electron microscopy images of representative sorted IgD-ME (top left) or IgG/A-ME (bottom left) B cells as well as a representative sorted IgD-PC (top right) or a representative sorted IgG/A-PC (bottom right). Red circles, perinuclear area; red arrowheads, Golgi structures; blue arrowheads, RER; scale bars in full cell images, 2 μm; scale bars within insets, 500 nm.

To verify whether these transcriptional changes corresponded to specific ultrastructural properties, we inspected the morphology of IgD-ME B cells and IgD-PCs by transmission electron microscopy (TEM). Similar to IgD-PCs but also IgG-PCs and IgA-PCs, IgD-ME B cells exhibited abundant rough endoplasmic reticulum (RER) and large perinuclear Golgi apparatus, although in a less pronounced manner (**Fig. 5D**). Together with functional and transcriptional data, these ultrastructural data suggest that IgD-ME B cells serve as a ready-to-differentiate reservoir of IgD-PCs for steady-state IgD responses.

### Tonsillar IgD-ME B cells constitute the preferential precursors of IgD-PCs

We next wondered whether the clonal history of IgD-ME B cells included traces of their function as IgD-PC precursors. To address this question, we built phylogenetic trees by assigning V_H_DJ_H_ clones into lineage clusters as detailed in Methods. Then, we grouped lineage trees according to their isotype composition and considered all groups containing at least five trees per donor, to ensure statistical robustness. In this manner, we obtained groups of lineage trees encompassing IgD alone, IgM alone, combined IgG-IgA or combined IgM-IgG-IgA. To characterize the developmental history of cells forming these trees, we calculated the normalized tree height, i.e., the distance from the germline to the furthest tip in the tree divided by the number of non-germline sequences per tree. We found that IgD lineage trees were higher than trees with IgM alone, both IgG and IgA, or a combination of IgM, IgG and IgA (**Fig. 6A**), suggesting that IgD class-switched B cells have a longer and more complex clonal history compared to other tonsillar B cells.

**Fig. 6.**
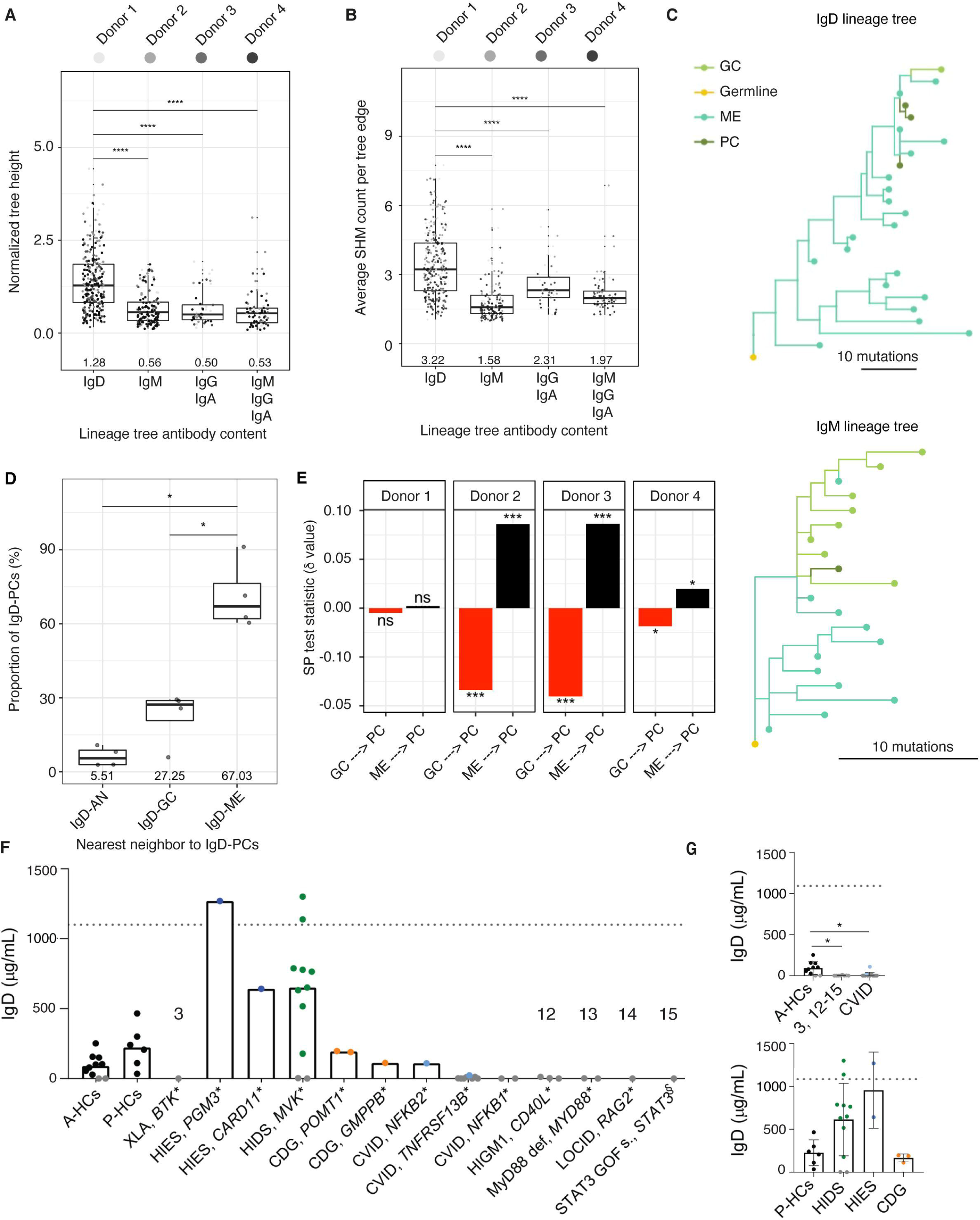
The IgD gene repertoire of tonsillar IgD class-switched cells exhibits a complex phylogenetic topology echoing the intricate *in vivo* requirements of the IgD response. (**A-B**) Phylogenetic topology analyses of lineage trees’ normalized heights (A) and average stepwise mutation count (B). (**C**) Examples of lineage trees obtained from the IgD repertoire (top) and IgM repertoire (bottom) of the same donor. The length of scale bars is equal to ten somatic hypermutations in both trees. (**D**) IgD-PC nearest neighbor analysis showing the proportion (%) of neighboring sequences to IgD-PC sequences. (**E)** Switch proportion (SP) test statistics values (δ) as performed on IgD lineage trees within each donor on transitions from IgD-GC B cells to IgD-PCs (GC→PC) and IgD-ME B cells to IgD-PCs (ME→PC). Numbers below plots in (A), (B) and (D) represent the median value per group. Statistical significance in (A), (B) and (D) was assessed with pairwise Mann-Whitney tests. ns, not significant; * p < 0.05, *** p < 0.001, **** p < 0.0001. (**F**) Total serum IgD measured by ELISA in adult healthy controls (A-HCs), pediatric healthy controls (P-HCs) as well as XLA, HIES, HIDS, FMF, CDG, CVID, HIGM1, MyD88 deficient, LOCID, or STAT3 gain-of-function syndrome (s.) patients. (**G**) Top panel. Total serum IgD in A-HCs, pooled CVID and pooled XLA (position 3 from F), HIGM1, MyD88 deficient, LOCID and STAT3 gain of function syndrome patients (positions 12-15 from F). Bottom panel. Total serum IgD in P-HCs, HIDS patients, pooled HIES patients, and pooled CDG patients. Gray symbols, below the limit of detection values; dotted line, saturation point of the assay; *, loss of function mutation; §, gain-of-function mutation. Differences were assessed with Kruskal-Wallis test followed by a post-hoc Dunn’s test. * p < 0.05. Statistical significance was only calculated between A-HC or P-HC and other patient cohorts. Comparisons lacking statistical reporting are not statistically significant.

Furthermore, we assessed the mutation activity in IgD class-switched lineage trees by calculating the average mutation count per tree edge. This analysis showed that IgD lineage trees had a higher average mutation count per tree edge compared to trees including IgM alone, IgG and IgA or IgM, IgG and IgA (**Fig. 6B**). As reflected by their smaller scale bar, IgD lineage trees accumulated mutations at a higher rate compared to IgM trees (**Fig. 6C, Fig. S4**). These results confirm that IgD class-switched B cells form large clonal families composed of GC B cells, ME B cells and PCs that do not further switch to downstream isotypes but rather engage in a strong mutational activity.

To identify the precursors of IgD-PCs, we converted IgD trees into pairwise distance matrices, selected the nearest (minimal distance) non-PC neighboring sequence for each PC-originated sequence, and calculated the average nearest neighbor proportion (%) for each donor. This analysis revealed that IgD-ME B cells were the nearest neighbors to IgD-PCs rather than IgD-GC B cells (**Fig. 6D**), suggesting that IgD-ME B cells represent the most frequent precursor of IgD-PCs. However, this trend could also be explained by higher mutation rates in GC B cells causing them to be further diverged from the germline than IgD-ME B cells and IgD-PC at the time of tonsillectomy. In parallel, a switch proportion (SP) test performed on IgD lineage trees showed that the transition from IgD-ME B cells to IgD-PCs had a significantly higher SP statistic (δ) value in three of four donors compared to the transition from IgD-GC B cells to IgD-PCs (**Fig. 6E**).

This analysis indicates that IgD class-switched B cells may enter tonsillar GCs upon encountering aerodigestive antigens. After differentiating into GC intermediates, IgD class-switched B cells intensively mutate and generate IgD-ME B cells and IgD-PCs. Following secondary antigen recognition, IgD-ME B cells may undergo extra-GC differentiation into IgD-PCs without accumulating additional mutations, or differentiate into IgD-GCs that “edit” their IgD gene repertoire by accumulating additional mutations.

### IgD responses involve both innate and adaptive signals, including T helper type-2 signals

Knowing that *in vitro* induced IgD secretion involves signals from cytokine receptors, BCR, CD40, TACI or TLR ligands (*3*, *7*), we explored the *in vivo* impact of these signals by interrogating serum IgD from patients with rare inborn errors of immunity caused by or associated with mutations of genes relevant to antibody production. Compared to most healthy controls, serum IgD was below the limit of detection in 1) an X-linked agammaglobulinemia (XLA) patient with hypomorphic mutation of the *BTK* gene encoding the BCR-associated BTK kinase (*27*); 2) common variable immunodeficiency (CVID) patients with mutations of *TNFRSF13B* or *NFKB1* genes encoding CSR- and PC-inducing receptor TACI (*28*) or CSR-inducing NF-κB p50 (*28*), respectively; 3) two hyper-IgM type 1 syndrome (HIGM1) patients with mutations of the *CD40L* gene encoding CSR-inducing CD40L (*29*); 4) two MyD88-deficient patients with mutations of the *MYD88* gene encoding the TLR adaptor MyD88 (*30*); 5) a late onset combined immunodeficiency (LOCID) patient with hypomorphic mutations of the *RAG2* gene encoding lymphopoiesis-orchestrating RAG2 (*27*, *31*); and 6) a STAT3 gain-of-function syndrome patient with a gain-of-function *STAT3* mutation abnormally increasing signals from lymphocyte-activating STAT3 (*27*) (**Fig. 6F**, **Table S1**). Compared to age-matched adult healthy controls, pooled CVID patients or pooled XLA, HIGM1, MyD88 deficiency, LOCID and STAT3 gain-of-function syndrome patients showed significantly reduced serum IgD (**Fig. 6G**).

In contrast, serum IgD was normal or tendentially increased in 1) two hyper-IgE syndrome (HIES) patients with immunodeficiency and pro-inflammatory T_H_2 cell responses linked to mutations of phosphoglucomutase 3-encoding *PGM3* or CARD11-encoding *CARD11* (*32*), with the *PGM3* mutation causing IgD secretion above the assay saturation limit; 2) several patients with hyper-IgD syndrome (HIDS), an autoinflammatory syndrome with mutation of *MVK* encoding mevalonate kinase (*33*); 3) two patients with congenital disorders of glycosylation (CDG) involving impaired O-glycosylation due to mutations of protein O-mannosyltransferase 1-encoding *POMT1* or GDP mannose pyrophosphorylase B-encoding *GMPBB* (*34*); and 4) a CVID patient with mutations of *NFKB2* encoding survival-inducing NF-κB p52 (*28*) (**Fig. 6, F** and **G**, **Table S1**). Thus, IgD responses may involve NF-κB p50-signals from BCR, CD40, TACI and TLR receptors as well as STAT3 signals from cytokine receptors and may increase in the presence of inflammation, including T_H_2 cell-driven inflammation.

### IgD responses target common nasopharyngeal antigens

To dissect the reactivity of IgD antibodies to common nasopharyngeal antigens, we generated a set of recombinant monoclonal antibodies (mAbs) from sorted tonsillar IgD-GC B cells. IgG mAbs from memory B cells specific to the receptor-binding domain of severe acute respiratory syndrome coronavirus-2 were expressed in parallel with IgD mAbs and used as controls. Briefly, IGHV and IGLV sequences from 20 single-sorted IgD-GC B cells with > 10 IGHV mutations, and 3 from IgD-GC B cells with ≤10 IGHV mutations (mAbs 3, 9 and 10) were expressed as IgG1 mAbs after sequencing (**Table S2**). Like control mAbs, mAbs from IgD-GC B cells encompassed Cγ1 to facilitate their downstream analysis (*5*). As shown by ELISA, some tonsillar IgD mAbs recognized milk, egg, pollen, dust mite, seaweed, viral and/or fungal antigens at greater levels than control mAbs (**Fig. 7A, Fig S5A**). As shown earlier (*5*), tonsillar IgD mAbs also bound self-antigens, including single-stranded DNA (ssDNA), double-stranded DNA (dsDNA), and/or insulin (**Fig. 7A, Fig S5A**). Of all IgD mAbs studied, mAbs 10, 47 and 48 showed higher polyreactivity with detectable binding to most nasopharyngeal antigens tested (**Fig. 7B**). As shown by flow cytometry, IgD mAbs also recognized isolated strains of aerodigestive bacteria, but did not show significant binding to intestinal bacteria, including *E. coli* (**Fig. 7, C** and **D, Fig. S5B**).

**Fig. 7.**
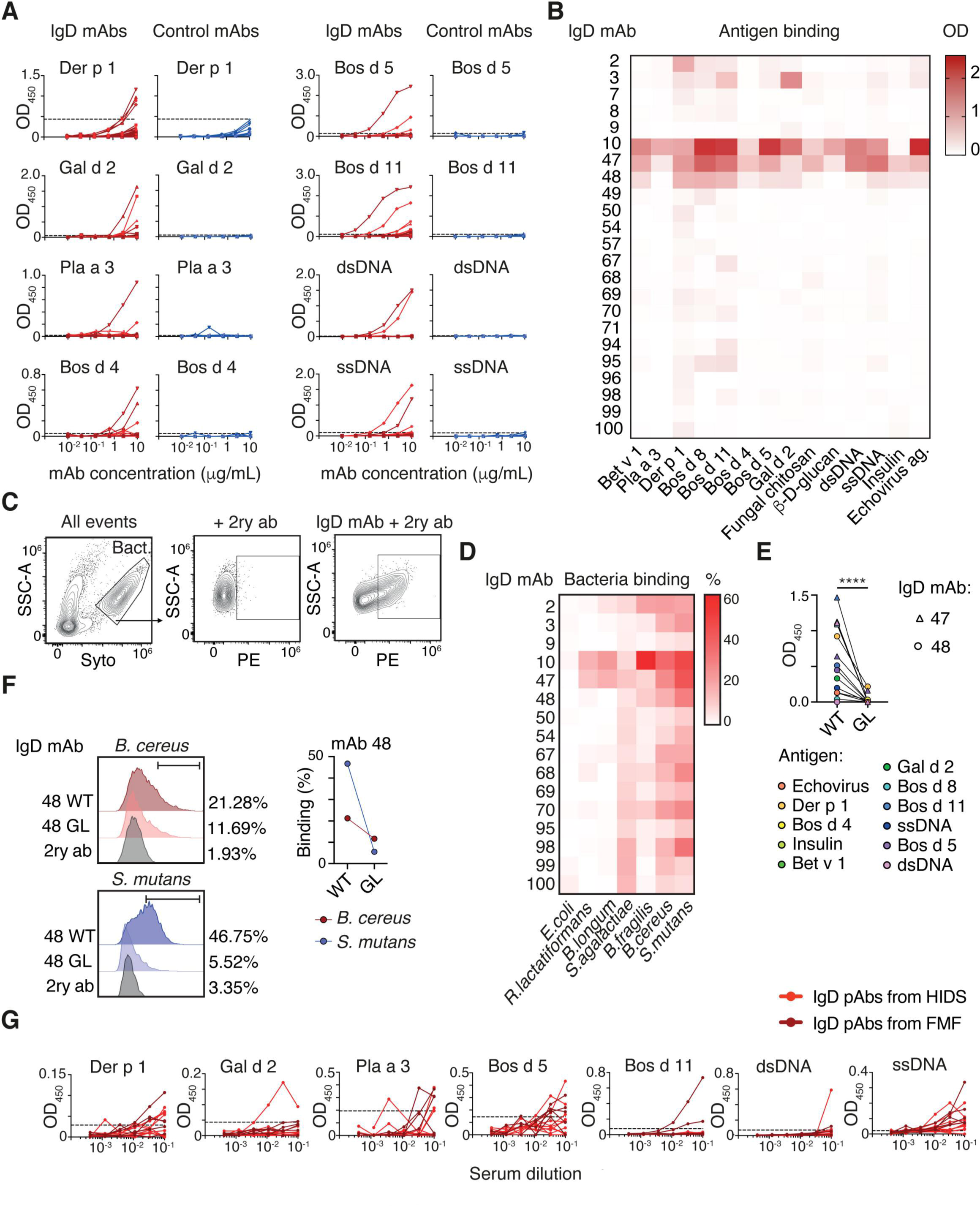
Tonsillar or systemic IgD antibodies broadly react against common aerodigestive antigens and oral commensal bacteria. (**A**) ELISA measuring the binding of 23 recombinant IgD mAbs from single-sorted tonsillar IgD class-switched GC B cells (red) to selected aerodigestive and autologous antigens. This binding was also measured in 10 control recombinant IgG, IgA or IgM mAbs from single-sorted circulating canonical memory B cells (blue) specific to the receptor-binding domain of the SARS-CoV-2 spike protein. All mAbs encompassed Cγ1, so that reactivity differences only stemmed from the antigen-binding variable region. Dashed lines indicate reactivity thresholds. (**B**) Heatmap summarizing binding intensity to antigens as in (A) by IgD mAbs at 10 μg/mL. (**C**) Representative flow cytometry gating of Syto^+^ bacteria and IgD-bound bacteria. (**D**) Heatmap summarizing percentage of selected bacterial isolates bound by IgD mAbs, determined as in (C). (**E**) ELISA binding of wild type (WT) or germline (GL) mAbs 47 (triangles) and mAb 48 (circles) against selected antigens from (A-B). (**F**) Representative histograms (left) and summary graph (right) of WT and GL mAb 48 binding to *B. cereus* and *S. mutans*. (**G**) Binding of circulating IgD pAbs from HIDS and familial Mediterranean fever (FMF) patients with reactive hyper-IgD production (n = 15) to antigens as in A. Dashed lines indicate reactivity thresholds. Significance was determined with a paired Wilcoxon test. ****p < 0.0001.

While the VH region of IgD mAbs 10 and 48 was encoded by IGHV3-30, the VH region of mAb 47 was encoded by IGHV4-34 (**Table S2**). Consistent with our earlier data, these IGHV genes are highly utilized by IgD-GC B cells, IgD-ME B cells and IgD-PCs and, as shown by others (*9*, *22*, *35*–*37*), encode highly polyreactive and autoreactive antibodies. Unlike IgD mAb 10, mAbs 47 and 48 were extensively mutated (**Fig. S5C**, **Table S2**), raising the possibility that extensive SHM allows some IgD-GC B cells to mitigate their autoreactivity while maintaining or even increasing their polyreactivity (*38*, *39*).

To evaluate the impact of SHM on the reactivity of IgD mAbs, we generated germline revertants of mAbs 47 and 48 by replacing their mutated bases with bases from the corresponding putative germline sequence (**Fig S5C**). Compared to wild-type controls, these germline revertants showed decreased reactivity to airborne, food, viral, and self-antigens (**Fig. 7E**). Similarly, the germline revertant of mAb 48 showed less reactivity to oral bacteria such as *Bacillus cereus* and *Streptococcus mutans* (**Fig. 7F**).

To evaluate the impact of the Cγ1 chain on the reactivity of recombinant IgD mAbs, we engineered two of these mAbs to encompass Cδ instead of Cγ1 and found comparable reactivity (**Fig. S5D**). To further confirm earlier reactivity results, we studied native IgD pAbs from the circulation. Given that secreted IgD is present at very low concentration in the serum from healthy individuals, we took advantage of serum from patients with HIDS and familial Mediterranean fever (FMF), two autoinflammatory disorders with reactive hyper-IgD production. Similar to recombinant tonsillar IgD mAbs, native serum IgD pAbs bound to common aerodigestive antigens (**Fig. 7G**). In summary, IgD-PCs may release polyreactive IgD to clear common nasopharyngeal antigens (**Fig. 8**). Due to their pre-plasmacellular properties, IgD-ME B cells would readily differentiate to IgD-PCs, thereby serving as a ready-to-use IgD-PC reservoir for homeostatic IgD responses. Aside from supporting a mutation-intensive pathway instrumental to enhance the polyreactivity of class-switched IgD, sustained exposure to nasopharyngeal antigens may account for the atypical properties of IgD-ME B cells.

**Fig. 8.**
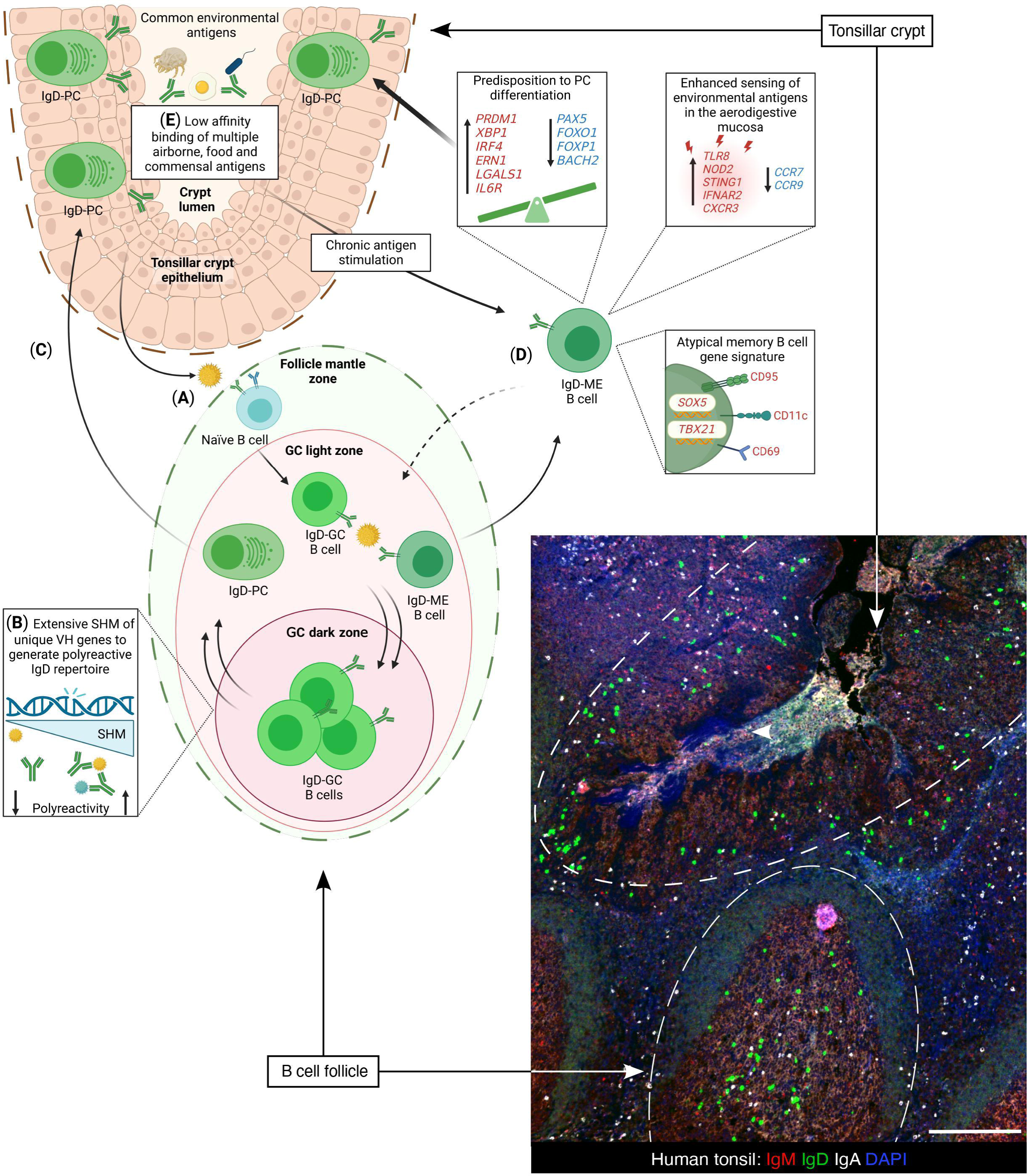
Tonsillar IgD-ME B cells form a ready-to-use pre-plasmacellular repertoire for steady-state IgD responses. (**A**) Highly conserved epitopes from common environmental antigens, including airborne, food and commensal antigens, select a unique tonsillar fraction of follicular naïve IgD^+^IgM^+^ B cells. The ensuing activation induces CSR from IgM to IgD. (**B**) The resulting IgD class-switched IgD^+^IgM^−^ B cells enter the GC and differentiate to IgD-GC B cells, which undergo extensive SHM through a program aimed at increasing IgD polyreactivity while attenuating IgD autoreactivity. (**C**) IgD-GC B cells clonally differentiate to IgD-PCs, which release IgD after migrating to nasopharyngeal effector sites, including the crypt epithelium, in response to chemotactic signals likely derived from CXCR3. (**D**) Alternatively, IgD-GC B cells clonally differentiate to IgD-ME B cells, which express higher *TBX21* and *SOX5* along with higher CD11c, CD69, CD95 and CXCR3. This atypical memory B cell signature likely results from persistent IgD-ME B cell exposure to aerodigestive antigens. Continuous antigen exposure could also account for the pronounced pre-plasmacellular properties of IgD-ME B cells, including higher *PRDM1*, *IRF4*, and *XBP1* expression combined with lower *PAX5*, *BACH2*, and *FOXP1* expression. Consistently, IgD-ME B cells readily differentiate to IgD-PCs *in vitro*. By serving as dominant IgD-PC precursors over IgD-PCs, IgD-ME B cells are key to mount IgD responses that are highly dependent *in vivo* on both innate and adaptive signals, including TLR signals. (**E**) By targeting common environmental antigens, IgD antibodies from IgD-PCs may mitigate the pro-inflammatory impact of these antigens. Figure created with BioRender.com.

## DISCUSSION

Here we have shown that human tonsils include atypical IgD^+^IgM^−^ B cells with unique BCR gene repertoire, complex developmental history, pronounced pre-plasmacellular properties, and extensive clonal relatedness to IgD-PCs. These IgD-ME B cells acquired reactivity to multiple common nasopharyngeal antigens through a mutation-dependent pathway that heightened IgD secretion in response to inflammatory signals, including pathological T_H_2 signals.

IgD-PCs were described more than two decades ago (*13*), but their ontogeny, regulation, clonal architecture, and reactivity to common nasopharyngeal antigens remain poorly characterized. Clarifying these properties may help understand the role of nasopharyngeal IgD responses in immune tolerance as well as inflammation (*40*, *41*). Besides broadening our understanding of humoral responses to potential allergens such as common food and airborne proteins (*42*), this advance could facilitate the development of novel prognostic tools and therapeutic strategies for allergic disorders. Indeed, allergic children with robust allergen-specific IgD responses show a lower risk of anaphylaxis and increased likelihood of naturally outgrowing allergy (*43*, *44*). Increased IgD responses to allergens are also associated with desensitization or even sustained tolerance in allergic patients treated with oral immunotherapy (*44–46*).

IgD could optimize antigen clearance through IgD-binding basophils and mast cells (*46–48*). In these cells, IgD ligation by antigen attenuates IgE-mediated degranulation but promotes IL-4, IL-5 and IL-13 release, which amplifies T_H_2 cell-mediated B cell production of antigen-specific IgG and IgE (*46*, *48*). Consistent with data recently published (*44–46*), secreted IgD may deploy its protective function as part of a non-inflammatory T_H_2 cell-mediated response aimed at promoting tolerance to common aerodigestive antigens, including potential allergens.

We furthered our understanding of IgD responses by identifying and characterizing a hitherto elusive tonsillar subset of highly mutated IgD-ME B cells sharing various properties with atypical ME B cells. IgD-ME B cells also exhibited clonal affiliation to IgD-GC B cells and IgD-PCs and displayed a complex developmental history compatible with their clonal origin from IgD-GC B cells and clonal differentiation to IgD-PCs. This last finding echoes recent studies showing that age-associated atypical memory B cells serve as progenitors of antibody-secreting plasmablasts (*49*).

Compared to canonical IgG-ME and IgA-ME B cells, IgD-ME B cells were enriched in gene products linked to key aspects of PC biology, including the UPR, protein trafficking from or to the RER and Golgi apparatus, and transport of protein-containing cargos. Aside from showing up-regulated gene PC identity products like *SDC1*, *ERN1*, *IRF4*, *BLIMP1*, *XBP1*, and *IL6R*, IgD-ME B cells displayed down-regulated B cell identity gene products like *PAX5*, *FOXO1*, *FOXP1*, and *BACH2*. By showing that IgD-ME B cells also displayed abundant RER and Golgi apparatus, our ultrastructural data suggest that these atypical ME B cells serve as a ready-to-differentiate reservoir of IgD-PCs for the initiation of rapid IgD responses.

Nasopharyngeal antigens could stimulate iterative entry of pre-existing hypermutated IgD-ME B cells into tonsillar GCs to elicit further IgD gene editing via SHM followed by IgD-PC differentiation. This process would permit IgD-ME B cells and their clonal IgD-PC progeny to continually adjust the reactivity of their BCRs to local antigen changes. IgD-ME B cells would further generate IgD-PCs through an extra-GC pathway that may afford more immediate protection. In general, IgD-ME B cells were found to retain a key property of ME B cells, i.e., the ability to rapidly become antibody-secreting PCs (*50*).

The widespread availability of IgD-reactive antigens is suggested by the clustering together of IGHV genes from tonsillar IgD-GC B cells, IgD-ME B cells and IgD-PCs from four different donors. The immunodominant nature of these antigens and their ability to bind intrinsically polyreactive IgD-BCRs would support the oligoclonal expansion of tonsillar IgD-GC B cells through a complex developmental pathway yielding IgD-ME B cells in addition to IgD-PCs. The involvement in this process of superantigen-binding sites from Cδ seems unlikely (*2*, *21*, *51*, *52*), as replacement mutations of IgD genes from IgD-GC B cells, IgD-ME B cells, and IgD-PCs preferentially targeted H-CDR rather than H-FRW regions, which suggests antigen-driven B cell selection in the GC (*19*, *20*). The little or no role of Cδ superantigen-binding sites was confirmed by the comparable reactivity of recombinant IgD mAbs expressing Cγ1 or Cδ. This reactivity was further validated in native IgD pAbs.

Consistent with earlier small-scale studies (*9*, *21*), all IgD class-switched B cell subsets, including IgD-ME B cells, were enriched in IGHV4-30, IGHV4-31, and, to a non-statistically significant degree, IGHV4-34. These IGHV genes are usually counter-selected by GCs due to their intrinsic autoreactivity (*37*). Their enrichment in IgD class-switched cells may reflect targeting by these cells of epitopes highly conserved across autologous and foreign nasopharyngeal antigens, including commensal antigens (*23*). Accordingly, IgD class-switched B cells from two donors also frequently used IGHV3-30, which encodes specificities targeting microbial polysaccharides (*53*). The concomitant reduced usage of IGHV3-13, IGHV3-48 and IGHV3-7 but also (although not statistically significant) IGHV3-23, which encode specificities targeting a broad range of antigens (*54*), could reflect the need by IgD class-switched B cells to focus their reactivity on a given set of nasopharyngeal antigens.

Some class-switched IgD antibodies recognized several airborne, food, fungal, bacterial, protist, and viral antigens, showing a reactivity that spanned across the known kingdoms of living things. This cross-kingdom polyreactivity could be aimed at continually and rapidly clearing common and generally harmless nasopharyngeal antigens in a non-inflammatory manner, possibly to enhance mucosal homeostasis. Consistent with earlier studies (*5*), the highly conserved nature of IgD-targeted epitopes would explain the autoreactivity of class-switched IgD antibodies, which may allow them to clear apoptotic autologous cells in addition to foreign antigens. Germline reversion attenuated the polyreactivity of two of our IgD mAbs, raising the possibility that SHM is part of a GC-driven nasopharyngeal program aimed at generating tolerogenic IgD specificities to multiple common environmental antigens.

In addition to adaptive BCR and CD40 signals, IgD-ME B cells may take advantage of TACI, TLR and other innate signals to differentiate into IgD-PCs (*3*, *7*). Consistently, IgD-ME B cells were enriched in *TNFRSF13B, TLR8* (encoding a surface receptor for ssRNA) as well as *STING2* (encoding an intracellular receptor for DNA) and *NOD1* (encoding an intracellular receptor for bacterial peptidoglycan) gene products. Accordingly, serum IgD was reduced in rare primary immunodeficiency disease patients with mutated *CD40L*, *TNFRSF13B*, *BTK* or *MYD88*. Being also enriched in *IL6R* and *IFNA2R*, IgD-ME B cells could further differentiate into IgD-PCs in response to STAT signals from IL-6 and IFN-α, which cooperatively induce PC differentiation (*26*). In this regard, a rare STAT3 GOF syndrome patient with mutated *STAT3* showed reduced serum IgD, although this reduction may result from an indirect mechanism causing lymphopenia and loss of memory B cells (*55*). The additional analysis of rare HIES patients with mutated *PGM3* and *CARD11* echoed published data showing that IgE-inducing T_H_2 cell-mediated signals, including IL-4, robustly enhance IgD responses (*46*). Similar to HIES, also HIDS cases with mutated *MVK* showed elevated serum IgD, which points to an enhancing impact of inflammation on the IgD response, a phenomenon also reported in autoimmunity (*7*). IgD secretion might compensatorily increase to enhance antigen clearance without aggravating inflammation due to the inability of IgD to recruit and activate complement (*40*).

The generation of IgD-PCs by IgD-ME B cells could further involve CXCR3, a CXCL10 receptor enriched on IgD-ME B cells compared to canonical IgG-ME and IgA-ME B cells. In the presence of IL-6, binding of CXCR3 by myeloid signals from CXCL10 enhances the differentiation of activated B cells into PCs (*7*). CXCR3 may also guide IgD-ME B cells from follicular inductive sites to CXCL10-expressing epithelial effector sites in the respiratory tract. The migration of IgD-ME B cells to the respiratory mucosa may be further facilitated by the down-regulation of the gut-homing receptors *CCR9* and *CCR10*. Chronic exposure of IgD-ME B cells to nasopharyngeal antigens may account for increased *ITGAX*, *FAS*, *CXCR3*, *SLAMF7*, *SOX5*, and *TBX21* gene expression, which is usually detected in mucosal ME B cells continually exposed to antigen (*56–58*).

A limitation of this study relates to the limited number of donors for high-throughput RNA-seq studies of tonsillar IgD-ME B cells, which was due to insufficient post-sorting cellular yield combined with insufficient RNA quality during the processing of some samples. This limitation was partly compensated by the high reproducibility and internal coherence of data from the available samples. An additional limitation relates to the limited power of studies involving primary immunodeficiency patients with known inborn errors of immunity, which are very rare. We attempted to compensate for this limitation by pooling together non-CVID samples from adult primary immunodeficiency donors with rare gene mutations. Finally, competition with abundant serum IgG pAbs reactive to common environmental antigens (*42*) likely limited the binding of serum IgD pAbs to the same antigens.

In summary, we have characterized a hitherto elusive subset of tonsillar IgD-ME B cells that serves as a ready-to-use IgD-PC reservoir for steady-state nasopharyngeal IgD responses. Considering recently published data (*43–46*), we propose that specificities resulting from IgD responses acquire polyreactivity to common aerodigestive antigens, including commensal, food and airborne antigens, via a mutation-dependent program designed to enhance mucosal tolerance.

## MATERIALS AND METHODS

### Study design

The primary aim of this study was to identify tonsillar IgD-ME B cells and characterize their phenotype, clonal architecture, transcriptional signatures, and reactivity properties. To address these questions and better understand the ontogeny and regulation of IgD responses, we combined standard and high-throughput approaches with sorting of 13 tonsillar B cell subsets, immunoglobulin gene repertoire analysis, cloning and expression of recombinant IgD mAbs, and reactivity analysis of these antibodies as well as native IgD pAbs from both healthy controls and patients with rare inborn errors of immunity. Indeed, biological specimens from these patients offer the unique opportunity of characterizing the overall *in vivo* impact of immunologically relevant molecules on a given human immune response. We also analyzed native circulating IgD pAbs from HIDS samples to validate reactivity data obtained using recombinant tonsillar mAbs from healthy individuals. Indeed, serum IgD from HIDS patients permits to overcome a major limitation of serum IgD from healthy controls, i.e., its scarce concentration. We first defined the topography, phenotype, transcriptional signatures, ultrastructural properties, and fundamental functions of tonsillar IgD-ME B cells as well as canonical IgG-ME and IgA-ME B cells by combining standard approaches with high-dimensional flow cytometry, RNA sequencing, and transmission electron microscopy. To further investigate the clonal architecture of IgD-ME B cells and their relationship with other tonsillar IgD class-switched B cell subsets, we performed high-throughput Ig gene sequencing, followed by clonal lineage and phylogeny analyses. Next, Ig cloning-expression technology, ELISA and bacterial flow cytometry were applied to assess the reactivity of secreted IgD to common nasopharyngeal antigens. Finally, total serum IgD from patients with rare inborn errors of immunity provided insights into the *in vivo* regulation of IgD responses. Experiments were performed 2-6 times at least in triplicate (where applicable) and showed excellent reproducibility.

### Tissue and blood samples

Tonsil samples were collected from donors tonsillectomized as a result of tonsillar hypertrophy. Heparinized blood samples were acquired from adult healthy donors. Splenic samples were obtained from deceased healthy donors or individuals subjected to post-traumatic splenectomy. The Ethical Committee for Clinical Investigation of the Institut Hospital del Mar d’Investigacions Mèdiques approved the use of blood and tissue samples (CEIC-IMIM 2011/4494/I, 2014/5892/I and 2022/10464/I). Fresh samples and formalin-fixed or paraffin-embedded tissue sections were obtained from the Mar Biobanc tissue repository with patient-signed informed consent. Archived patient serum samples from Hospital Sant Joan de Déu (Barcelona), Hospital Clinic (Barcelona) and The Mount Sinai Hospital (New York) were obtained from their respective biobanks with patient-signed informed consent.

### Sample processing

Tonsillar mononuclear cells were obtained by perfusion of fresh tissue specimens with sterile phosphate buffer solution (PBS) (Biowest). Peripheral blood mononuclear cells (PBMCs) were isolated from heparinized blood samples by separation on a Ficoll-Hypaque gradient (GE Healthcare). For detailed splenocyte isolation protocol, see Supplementary Materials and Methods.

### Cell flow cytometry

Single-cell suspensions of tonsillar, splenic or peripheral blood mononuclear cells were stained with primary antibodies using the same protocol (**Table S3)**, which is detailed in Supplementary Materials and Methods. All flow cytometry data were analyzed using FlowJo V10 software (TreeStar). T-distributed stochastic neighbor embedding (t-SNE) projections from spectral flow cytometry data were generated using the default settings on FlowJo’s build-in t-SNE function, and based on the surface expression of CD10, CD11c, CD21, CD27, CD38, CD43, CD69, CD95, CXCR3, and Igλ light chain. Then, clusters were defined using the XShift clustering algorithm (*59*) and projected onto t-SNE plots for visualization.

### FACSorting

A detailed protocol is provided in Supplementary Materials and Methods. Discrete B cell or PC populations were selected according to forward scatter (FSC) and side scatter (SSC) parameters as well as expression of specific surface molecules (**Fig. S1**). Only IgA-ME and IgG-ME B cells expressing CD27 were included in functional, transcriptional and TEM studies. Repertoire and phenotypic studies included both dominant CD27^+^ and small CD27^−^ fractions of IgA-ME and IgG-ME B cells (*57*, *60*).

### B cell cultures

Gated tonsillar naïve, IgM-ME, IgD-ME, IgA-ME and IgG-ME B cells (**Fig. S1)** were sorted and processed as described in Supplementary Materials and Methods. Culture supernatants were saved for Ig quantification by ELISA and differentiation was assessed by flow cytometry.

### Immunofluorescence staining and image processing

Formalin-fixed paraffin-embedded 3-μm thick human tonsil sections were processed as detailed in Supplementary Materials and Methods. Samples were stained with various combinations of primary antibodies (**Table S4**) for 2 h at room temperature. After washing, fluorochrome-conjugated secondary antibodies (**Table S4**) were added to the tissue together with DAPI and incubated for 1 h at room temperature. After washing, FluorSave Reagent (Millipore) was applied and coverslips were fixed with CoverGrip Coverslip Sealant (Biotium). Images were acquired using a Nikon Eclipse Ni-E microscope and processed with ImageJ software.

### Recombinant monoclonal antibody production and purification

Plasmids encoding human IgD mAbs (*5*) were provided by Patrick C. Wilson and processed according to a mAb production and purification protocol detailed in Supplementary Materials and Methods. Peripheral blood IgG, IgA and IgM mAbs specific for the receptor-binding domain of severe acute respiratory syndrome coronavirus-2 were expressed in parallel with tonsillar IgD mAbs as described earlier (*61*) and used as control mAbs. Of note, all recombinant mAbs were synthesized to express the Fc domain from IgG1, i.e., Cγ1, which facilitated their purification. However, each mAb express the original antigen-binding Fab domain.

### Generation of germline revertants

Mutations in the germline sequences of IGHV and IGLV of mAb 47 and mAb 48 were determined using IgBlast. Full-length IGHV-D genes, including CDR3 regions, were identified following the criteria of Corbett et al. (*62*). The germline sequences were ordered as gblocks from IDT. The vectors were linearized with EcoRI-HF and Sal-HF or EcoRI-HF and XhoI restriction enzymes for the heavy and light chains, respectively. Linearized vectors were purified from gel and respective germline sequences were cloned using Gibson assembly method (*63*). New constructs were verified by Sanger sequencing, and germline revertant mAbs were produced and purified as described above.

### IgD ELISA

Total IgD from cell culture supernatants was measured using the Human IgD ELISA Quantitation Set (Bethyl Laboratories) according to the manufacturer’s instructions. A detailed protocol is described in Supplementary Materials and Methods. Total IgD from serum was measured using the IgD Human ELISA Kit (Abnova) according to the manufacturer’s instructions. In both assays, concentrations were calculated by extrapolating sample absorbance values with values from the standard curve.

### IgM, IgA and IgG ELISAs

Concentrations of IgM, IgG and IgA in cell culture supernatants were measured as detailed in Supplementary Materials and Methods.

### Antibody reactivity analysis

The reactivity of both recombinant and serum antibodies was determined by ELISA as detailed in Supplementary Materials and Methods using commercially available antigens (**Table S5**).

### Bacterial reactivity flow cytometry

Bacterial isolates of *Escherichia coli* (ATCC, 25992), *Bacteroidetes fragilis* (ATCC, 25285), *Bifidobacterium longum* (ATCC, 15707), *Ruthenibacterium lactatiformans* (ATCC, 100348), *Bacillus cereus* (ATCC, 11778), *Streptococcus mutans* (ATCC, 25175) and *Streptococcus agalactiae* (ATCC, 13813) were heat-inactivated at 65°C for 20 min. A detailed flow cytometry protocol is provided in Supplementary Materials and Methods.

### RNA-seq

Tonsillar cells were FACSorted as described above. Cells were centrifuged and homogenized using QIAshredder (Qiagen). RNA was isolated with RNeasy Micro Kit (Qiagen) according to manufacturer’s instructions. RNA quality and quantity were assessed by Bioanalyzer using Agilent RNA 6000 Pico Kit (Agilent Technologies). Samples used for analysis had RIN ≥ 6.4. NGS libraries with polyA capture were prepared according to the protocol of NEBNext® Ultra™ II detailed in Supplementary Materials and Methods.

### Bioinformatic analysis of transcriptomic data

Raw sequencing reads in the fastq files were mapped with STAR version 2.7.8a (*64*) Gencode release 41 based on the GRCh38.p13 reference genome and the corresponding GTF file. The table of counts was obtained with featureCounts function in the package subread, version 2.0.3 (*65*). Filtering of lowly expressed genes was done by keeping genes having more than 1 CPM in at least 4 samples. Raw library size differences between samples were treated with the weighted “trimmed mean method” TMM (*66*) implemented in the edgeR package (version 3.36.0). The normalized counts were used in order to make unsupervised analysis, PCA and clusters. Genes starting with IgH (except for IGHMBP2), IgK or IgL (except for those starting with IgLL and IgLon5) were excluded. For the differential expression (DE) analysis, read counts were converted to log2-counts-per-million (logCPM) and the mean-variance relationship was modeled with precision weights using voom approach in limma package. SVA package (version 3.38.0) was used to compute surrogate variables (SV) using svaseq function and all SVs were added to the design matrix with the primary variable. Donor variable was also included in the model. FDR was used to adjust for multiple comparisons. Genes were considered differentially expressed if adj.P.Value < 0.05 and |log2FC| > 1. All analyses were done with R version 4.1.2. Pre-Ranked Gene Set Enrichment Analysis (GSEA) (*67*) implemented in clusterProfiler package (version 4.0.0) (*68*) was used to retrieve enriched functional pathways (Reactome from MsigDB, version 7.5.1). The ranked list of genes was generated using the - log(p.val)*signFC for each gene from the statistics obtained in the DE analysis. Gene lists for heatmaps manually curated based on the Panther v17.0 classification (*69*) of the differentially-expressed gene lists.

### Transmission electron microscopy (TEM)

Tonsillar B cells and PCs were sorted as described earlier and processed according to a protocol detailed in Supplementary Materials and Methods.

### Sequencing of Ig gene repertoire

Sorted cells were lysed with Qiashredder columns (Qiagen) following the manufacturer’s protocol. Cellular RNA was isolated from sorted B cell subsets with RNeasy Micro Kit (Qiagen) following the manufacturer’s instructions, with a final elution in 14 μL RNAse-free water. 10 μL RNA were used for reverse transcription into cDNA using TaqMan® Reverse Transcription Reagents with random hexamers (Thermo Fisher). For IGHV gene PCR, 5 μL of cDNA were mixed with High-Fidelity Platinum PCR Supermix (Thermo Fisher) containing 50 nM forward primers specific for the framework region 1 of VH1, VH2, VH3, VH4, VH5 or VH6 and 250 nM primers specific for Cδ, Cμ, Cγ or Cα and encompassing corresponding Illumina Nextera sequencing tags along with unique molecular identifiers (UMI) (**Table S6**). For this study, UMI-based correction was not performed (*70*, *71*). Further amplification and sequencing protocols are described in detail in Supplementary Materials and Methods. In total, 120767500 IGHV gene sequences from 4 donors were obtained through NGS.

### Preprocessing of Ig gene repertoire sequencing

Raw sequencing reads for each sample from the two MiSeq runs were merged and preprocessed as detailed in Supplementary Materials and Methods.

### Clonal lineage clustering and lineage tree construction

We used the Change-O toolkit to parse the preprocessed dataset in blast format using the command MakeDb.py (*70*). Subsequently, we assigned our clones into clusters using the hierarchicalClones command from the SCOPer package (version 1.1) (*71*). Sequences from the same donor with identical V and J germline genes and 85% junction nucleotide sequence similarity were assigned to a single lineage cluster. Maximum parsimony phylogenetic trees were built from clonal lineages including 20-200 clonal sequences per lineage using the R package dowser (version 0.0.4) (*72*).

### Phylogenetic topology analyses

To perform nearest neighbor analysis, we included only lineage trees that exclusively contained IgD sequences, for a total of 32, 59, 82 and 101 clonal lineages (i.e., trees) for donors 1–4, respectively. Further analysis is detailed in Supplementary Materials and Methods.

### Trait-phylogeny association analysis

For this analysis, we included clonal lineages that exclusively contained IgD sequences. We firstly examined the association significance between tree trait values (cell types) and tree topology using the parsimony score test (PS test), as described previously (*72*) and further detailed in Supplementary Materials and Methods.

### Somatic hypermutation analysis

We used the SHazaM R package (version 1.0.2) to compute the count and frequency of V gene replacement (R) and silent (S) mutations (*70*). Further details are provided in Supplementary Materials and Methods.

### Quantification of clonal persistence (overlap)

A clone was defined as a unique V Gene-CDR3 (aa sequence)-J Gene. Pairwise clonal persistence (CP) between repertoires A and B was calculated as follows (*73*):

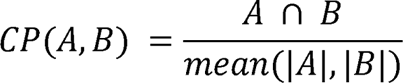

Where A ∩ B is the number of non-redundant shared clones between A and B, while |A| and |B| refer to the number of unique clones in repertoires A and B, respectively.

### V gene usage and heatmap generation

Germline V gene repertoires for each sample were calculated by determining the frequency of VDJ clones that map to a particular V-gene independently of the clone occurrence frequency in the repertoire (*74*). We used the ComplexHeatmap R package (version 2.5.1) to generate our heatmap following a complete-linkage hierarchical clustering algorithm, where pairwise distances among rows and among columns were calculated by Euclidean distance (*75*).

### Statistics and graphical visualization

The Ig gene repertoire analysis, RNA-seq statistical analysis and graphics generation were performed in RStudio (R version 4.0.3) (*76*). We used packages ggpubr, version 0.4.0 (*77*), and rstatix, version 0.6.0 (*78*) for statistical testing as well as packages ggplot2, version 3.3.3 (*79*), and cowplot, version 1.1.1 (*80*), for data visualization, unless stated otherwise. For phenotypic and functional studies, statistical analysis was performed using Prism 5.03 software (GraphPad). Details regarding statistical testing are indicated in the figure legends.

## Acknowledgments

We thank Patrick C. Wilson for providing plasmids encoding tonsillar IgD mAbs; Kenneth Hoehn for helping with the assembly of phylogenetic tree analysis; personnel from the flow cytometry facility at PRBB for providing technical support and helping with cell sorting; personnel from the genomics facility at PRBB for helping with experimental design and DNA sequencing; personnel from the Pathology Department of Hospital del Mar for providing tonsillar samples; personnel from MARGenomics service at PRBB for providing excellent technical support and guidance; personnel from the electron microscopy service at UAB for providing excellent technical support. We are indebted to the “Biobanc de l’Hospital Infantil Sant Joan de Déu per a la Investigació” integrated in the Spanish Biobank Network of ISCIII for the sample and data procurement. We also thank Mr. Javier Antunez-Sanchez (School of Life Sciences, the University of Warwick) for his help in BCR-seq data analysis. Finally, we are grateful to Kang Chen for reading the manuscript and Daniel Garzón Segura for helping with cellular immunology techniques.

## Funding

Ministerio de Ciencia, Innovación y Universidades grant RTI2018-093894-B-I00 (AC) Ministerio de Ciencia, Innovación y Universidades grant SAF2014-52483R (AC) European Research Council, European Advanced Grant ERC-2011-ADG-20110310 (AC) Institute of Health Carlos III, Miguel Servet Research Program MS19/00002 (GM) Ministerio de Ciencia e Innovación, Ramon y Cajal grant RYC2021-031642-I (GM) UiO World-Leading Research Community, UiO:LifeSciences Convergence Environment Immunolingo, EU Horizon 2020 iReceptorplus #825821 (VG)

Research Council of Norway project #300740 (VG)

Research Council of Norway project #331890 (VG)

## Author contributions

Conceptualization: AC, VG

Methodology: RT-P, HB, MF, ST-V, LC-M, GM, XM-F, JC, VG, AC

Formal analysis: RT-P, HB, MF, JP-B, JD-B

Investigation: RT-P, HB, MF, ST-V, LC-M, AS-G, JP-B, MG, XM-F, PC-H, JD-B, BA-R, MLR

Resources: LA, AG-G, AN-O, MP, JA, CC-R, JC, AC

Visualization: RT-P, HB, JP-B, JD-B, MF

Software: HB, AS, MC, VG

Data curation: RP-P, HB, MF

Funding acquisition: AC, VG, GM

Project administration: AC, RT-P

Supervision: AC, VG, GM

Writing – original draft: AC

Writing – review & editing: AC, VG, RT-P, HB, JG-M, MF, GM, SM

## Data and materials availability

All raw and processed RNA-seq data, as well as VDJ-seq data are deposited in publicly available databases. Correspondence and requests for materials should be addressed to A.C. and V.G.

## Supplementary Material

### Supplementary Materials and Methods

#### Splenocyte isolation

Fresh spleen samples were enzymatically digested for 40 min at 37°C in Hank’s balanced salt solution (Lonza) supplemented with 1 mg/mL collagenase IV (Thermo Fisher), 50 ng/mL DNase (New England Biolabs), and 0.5% human serum (Sigma-Aldrich). Splenic mononuclear cells were separated on a Ficoll-Hypaque gradient. All mononuclear cell suspensions were aliquoted and cryopreserved in fetal bovine serum (FBS) (Gibco) supplemented with 10% dimethyl sulfoxide (Merck) until the time of analysis.

#### Flow cytometry

Cells were incubated at 4°C in sterile staining buffer comprised of PBS at pH 7.4 supplemented with 5 g/L bovine serum albumin (BSA) (Sigma-Aldrich), 2 mM ethylenediaminetetraacetic acid (Merck), and 2 μL Fc-receptor blocking reagent (Miltenyi) for every 10^7^ cells. After 10-min incubation, cell suspensions were stained at 4°C for 30 min with a conventional or spectral B cell and plasma cell antibody cocktail (**Table S3**). For conventional flow cytometry, 4’-6-diamindine-20-phenylindole (DAPI) (Sigma-Aldrich) was used to exclude dead cells, and cells were acquired with a BD LSR Fortessa Cell Analyzer (BD Biosciences). In spectral flow cytometry, cells were acquired with an Aurora Cell Analyzer (Cytek). Dead cells were excluded using the LIVE/DEAD™ Fixable Yellow Dead Cell Stain Kit (Invitrogen).

#### FACSorting

Tonsil mononuclear cell suspensions were incubated for 10 min at 4°C with 2 μL Fc-blocking reagent for every 10^7^ cells in sterile staining buffer. Cells were then stained with a cell sorting-specific antibody cocktail (**Table S3**). Cell suspensions were filtered through a 35-µm cell strainer, prior to sorting with a FACSAria II (BD Biosciences). DAPI was used to exclude dead cells. B cells were sorted into tubes containing complete RPMI 1640 (Biowest), and immediately used for downstream analysis.

#### B cell cultures

FASCorted tonsiliar B cells were seeded (5 × 10^4^/well) in U-bottomed 96-well plates (Corning), and cultured for 6 days in complete RPMI 1640 supplemented with 10% FBS, 10 U/mL penicillin, 10 U/mL streptomycin (Biowest), and 2 mM L-glutamine (Biowest) with or without 100 ng/ml megaCD40L (Enzo Life Science) and 500 ng/ml IL-21 (Peprotech). Cells were microscopically monitored daily and counted using a Neubauer chamber. At day 6, well contents were transferred into new tubes and centrifuged at 300 g for 5 min.

#### Immunofluorescence staining

Formalin-fixed and paraffin-embedded 3-μm thick human tonsil sections were dewaxed and rehydrated by overnight incubation at 60°C, followed by treatment with a decreasing alcohol gradient. Heat-induced epitope retrieval was performed for 20 min in Tris-EDTA buffer (pH 9) and citrate buffer (pH 6). Sections were permeabilized by incubation with 0.2% Triton X (Merck) in PBS for 12 min. and blocked with 5% BSA (Miltenyi) or 5% Fc-receptor blocking reagent for 1 h at room temperature.

#### Recombinant monoclonal antibody production and purification

Antigen-binding Ig variable genes from both IgH and IgL chains of single-sorted tonsillar IgD^+^IgM^−^CD38^+^ B cells were amplified by RT-PCR and then cloned into Cγ1-encoding expression vectors. 150 × 10^6^ Expi293 cells seeded in 50 mL of Expi293™ Expression Medium (Thermo Fisher Scientific) were transiently transfected with 1 µg/mL of antibody-encoding plasmid DNA. After 5 days, cells were harvested by centrifugation for 5 min at 4000 rpm. The resulting supernatant was ultracentrifuged for 30 min at 30,000 rpm and filtered with a 0.45-µm Nalgene nylon filter (Thermo Scientific). Next, the filtered sample was applied onto a PBS-equilibrated HiTrap Protein A MabSelect or HiTrap LambdaMabSelect SuRe purification column (Cytiva) to purify Cγ1-or Cδ-expressing mAbs, respectively. The column was washed with 10 CV PBS and eluted with 100 mM glycine buffer, pH 3. The eluted sample was further subjected to glycine-to-PBS buffer exchange and concentrated using Amicon Ultra 50K filter. Purified mAbs and the corresponding flow through controls were loaded onto an SDS-PAGE gel for IgH and IgL bands visualization and quality control.

#### IgD ELISA

Human IgD ELISA Quantitation Set (Bethyl Laboratories) was used according to manufacturer’s instructions. Briefly, 96-well flat-bottomed plates (Nunc) were coated for 1 h with 10 μg/mL goat anti-human IgD coating antibody in a coating buffer comprised of 0.05 M carbonate-bicarbonate in PBS, pH 9.6. Wells were washed with a washing buffer comprised of ultrapure water supplemented with 50 mM Tris, 0.14 M NaCl, and 0.05 % Tween 20 at pH 8. Wells were blocked for 30 min at room temperature with a blocking buffer comprised of ultrapure water supplemented with 50 mM Tris, 0.14 M NaCl, and 1 % BSA at pH 8. Both samples and standards were diluted in dilution buffer comprised of ultrapure water supplemented with 50 mM Tris, 0.14 M NaCl, 1 % BSA and 0.05 % Tween 20 and incubated for 1 h at room temperature. After washing, a horseradish peroxidase (HRP)-labeled anti-human IgD antibody was added at 13.3 ng/mL and incubated for 1 h at room temperature. Plates were washed and developed using the TMB Substrate Reagent Set (BD Bioscience). Absorbance was read at 450 nm with an Infinite 200 PRO plate reader (Tecan).

#### IgM, IgA and IgG ELISAs

96-well flat-bottomed plates were coated with 1 μg/mL of goat anti-human Ig (Southern Biotech) in a coating buffer and incubated overnight at 4°C. After washing with PBS supplemented with 0.05% Tween 20 (PBS-T), wells were blocked with 1% BSA in PBS for 2 h at room temperature. Samples were diluted in PBS with 1% BSA and 0.05% Tween 20. A standard curve was prepared using purified human IgM (Calbiochem), IgA (Mpbio) and IgG (Mpbio) at 250 ng/mL and six 1:2 serial dilutions were performed. Both samples and standards were incubated for 2 h at room temperature. After washing, HRP-conjugated goat anti-human IgM (Southern Biotech), IgA (Southern Biotech) or IgG (Cappel) antibodies were added and incubated for 45 min. at room temperature. Plates were washed and developed using the TMB Substrate Reagent Set and absorbance was read at 450 nm with an Infinite 200 PRO plate reader (Tecan). Concentrations were calculated by extrapolating sample absorbance values with values from the standard curve.

#### ELISAs for antigen reactivity analysis

To screen for the reactivity of expressed monoclonal antibodies and serum samples, 96-well half-area flat bottom high-bind microplates (Corning) were coated overnight at 4°C with a given antigen (**Table S5**) diluted in PBS or with PBS alone for background subtraction. To measure Ig reactivity to insulin or chitosan, antigen was diluted in 1% acetic acid and neutralized with equimolar concentration of NaOH before coating. After washing with PBS-T, wells were blocked for 2 h at room temperature with PBS supplemented with 5% BSA and 0.1% Tween. To measure Ig reactivity to dsDNA, ssDNA or echovirus antigen, plates were blocked with PBS supplemented with 2% BSA. Recombinant mAbs were diluted to a concentration of 10 µg/mL in PBS supplemented with 1% BSA and 0.05% Tween 20. Serum samples were serially diluted starting with a 1:10 dilution. All samples were serially diluted 1:4 six times, added to the antigen-or PBS-coated plate, and incubated 2 h at room temperature. After washing, plates were incubated with HRP-conjugated anti-human IgG to detect mAbs, HRP-conjugated anti-human IgD to detect serum IgD, or HRP-conjugated anti-human Igλ (Southern Biotech) to compare the reactivity of Cγ1-expressing vs Cδ-expressing mAbs for 45 minutes at toom temperature. Plates were washed with PBS-T and developed with the TMB substrate reagent set (BD Biosciences). The development reaction was stopped with 1M H_2_SO_4_. Absorbance was read at 450 nm with an Infinite 200 PRO plate reader (Tecan). To quantitate the level of each antigen, optical density (OD) values were calculated after subtraction of the background defined as OD_450_ value of corresponding sample dilution on plates coated with PBS but no antigen. All negative values were normalized to 0. Threshold values were determined by calculating the mean ± 2 SD of the highest concentration OD_450_ value of all anti-RBD antibodies or control serum samples.

#### Bacterial reactivity flow cytometry

Heat inactivated bacterial isolates were incubated with recombinant mAbs for 30 min at room pemperature. After washing, bacterial pellets were incubated for 15 min at room temperature with polyclonal phycoerythrin (PE)-conjugated anti-human IgG antibodies (Infrared Laboratories). Bacteria incubated only with secondary antibodies were used as negative control to set up the PE-positive gate. Finally, bacterial samples were washed and resuspended in PBS with SYTO BC (Thermo Fisher; 1:60000). Contamination was minimized by passing all buffers and reagents through sterile 0.22 μm filters before use. Cells were analyzed using a Cytek Aurora Cytometer (Cytek Bioscience) with low FSC and SSC thresholds to allow bacterial detection. FSC, SSC and SYTO BC were set to a biexponential scale and samples were gated as SSC^+^SYTO BC^+^ and then assessed for PE-positive counts.

#### Transmission electron microscopy (TEM)

Cells were fixed with 2.5% glutaraldehyde in phosphate buffer 0.1 M for 2 h at 4°C, post-fixed with 1% osmium tetroxide with 0.8% potassium ferrocyanide for 2 h, and dehydrated with increasing concentrations of ethanol. Then, pellets were embedded in EPON resin (EMS, Hatfield) and polymerized at 60°C for 48 h. Sections of 70 nm in thickness were obtained with a Leica EM UC6 microtome (Wetzlar), stained with 2% uranyl acetate and Reynold’s solution consisting of 0.2% sodium citrate and 0.2% lead nitrate, viewed by a JEM-1400 transmission electron microscope (JEOL), and imaged at 120 kV voltage.

#### Sequencing of Ig gene repertoire

Amplification was performed with an initial step at 95 °C for 3 min., followed by 35 cycles at 95 °C for 30 sec., 58 °C for 30 sec., and 72 °C for 30 sec., supplemented with a final extension step of 72 °C for 5 min. PCR products were purified with AMPure XP beads (Beckman Coulter) and Nextera XT indices were added by PCR under the following conditions: 98°C for 30 sec., 5 cycles at 98°C for 10 sec., 63°C for 30 sec., and 72°C for 3 min. AMPure XP beads (Beckman Coulter) were used to purify PCR products, which were subsequently validated and pooled. The final pool was quantified by qPCR. Single-strand products were paired-end sequenced twice on a MiSeq instrument (Illumina) with the 600 Cycle v2 Kit (2 x 300 bp).

#### Preprocessing of Ig gene repertoire sequencing

The mean base call Phred quality score of all samples was ≥ 30. Subsequently, VDJ alignment and clonotyping were performed using MiXCR software package, version 3.0.12, considering the full VDJ region nucleotide sequence for clonotype assembly (*81*). In this way, a clone is defined as a unique VDJ region nucleotide sequence. We excluded clones containing stop codons and out-of-frame sequences. As described previously (*74*), we retained only those clones with at least a read count of two and with a CDR3 amino acid sequence length of more than three amino acids. For IgD class-switched IgD^+^IgM^-^ B cells, we only included IgD-unique clones for each sample by excluding IgD clones that have identical CDR3 amino acid sequence with IgM clones found in the same sample when sequenced with IgM-specific primers. Moreover, for each sample, we excluded those clones where the MiXCR-annotated constant gene did not match the reverse primer used to generate the sample (for example, IgM-annotated sequences in a sample where IgG primers were used).

#### Phylogenetic topology analyses

IgD-only trees were converted into pairwise distance matrices using the R package ape (version 5.4-1) (*82*) and we selected the nearest non-PC neighboring sequence for each PC-derived sequence (i.e., minimal distance). We calculated the proportion of neighboring sequences to IgD-PC sequences (%). For the normalized tree height analysis, pairwise distance matrices were used to extract tree height (i.e., distance from the germline to the furthest tip in the tree) and size (i.e., number of non-germline sequences per tree) information. Subsequently, we divided the tree height by the tree size to calculate the normalized tree height. For the average SHM count per tree edge analysis, we extracted pairwise tree edge lengths for each node and its parent from phylogenetic tree objects and calculated the corresponding average. In both normalized height and average SHM count per tree edge analyses, we included phylogenetic trees belonging to one isotype group that contained more than one cell type and had an occurrence of at least five trees per donor to ensure statistical robustness.

#### Trait-phylogeny association analysis

We calculated the significance of the PS test statistic using a permutation test, where the test statistic was recomputed for 1000 randomizations of trait values at the tips for each tree. We observed a significant PS test statistic in phylogenetic trees from all donors of the study. Following that, we used a restricted switch proportion test (SP test) as described previously (*72*) to examine the most likely ancestor to IgD-PC B cells across all trees for each donor. Similarly, using a permutation test, we calculated the significance of the SP test statistic for our phylogenetic trees within each donor by comparing observed test statistics to those obtained from 1000 randomizations of trait values at the tips for each tree. Of note, PS and SP test statistics were considered significant when accompanied with p-values < 0.05.

#### Somatic hypermutation analysis

We considered a VDJ clone to be mutated when harboring more than two mutations since the PCR step might be responsible for 1-2 mutations per clone (*83*). To compute the replacement:silent (R:S) ratio in mutated clones, we divided the number of R mutations by the number of S mutations. In sequences with R > 0 but no S mutations (S = 0), we followed an approach described previously (*84*), where the number of S mutations was set to one to avoid mathematically undefined results.

---

**Fig. S1.**
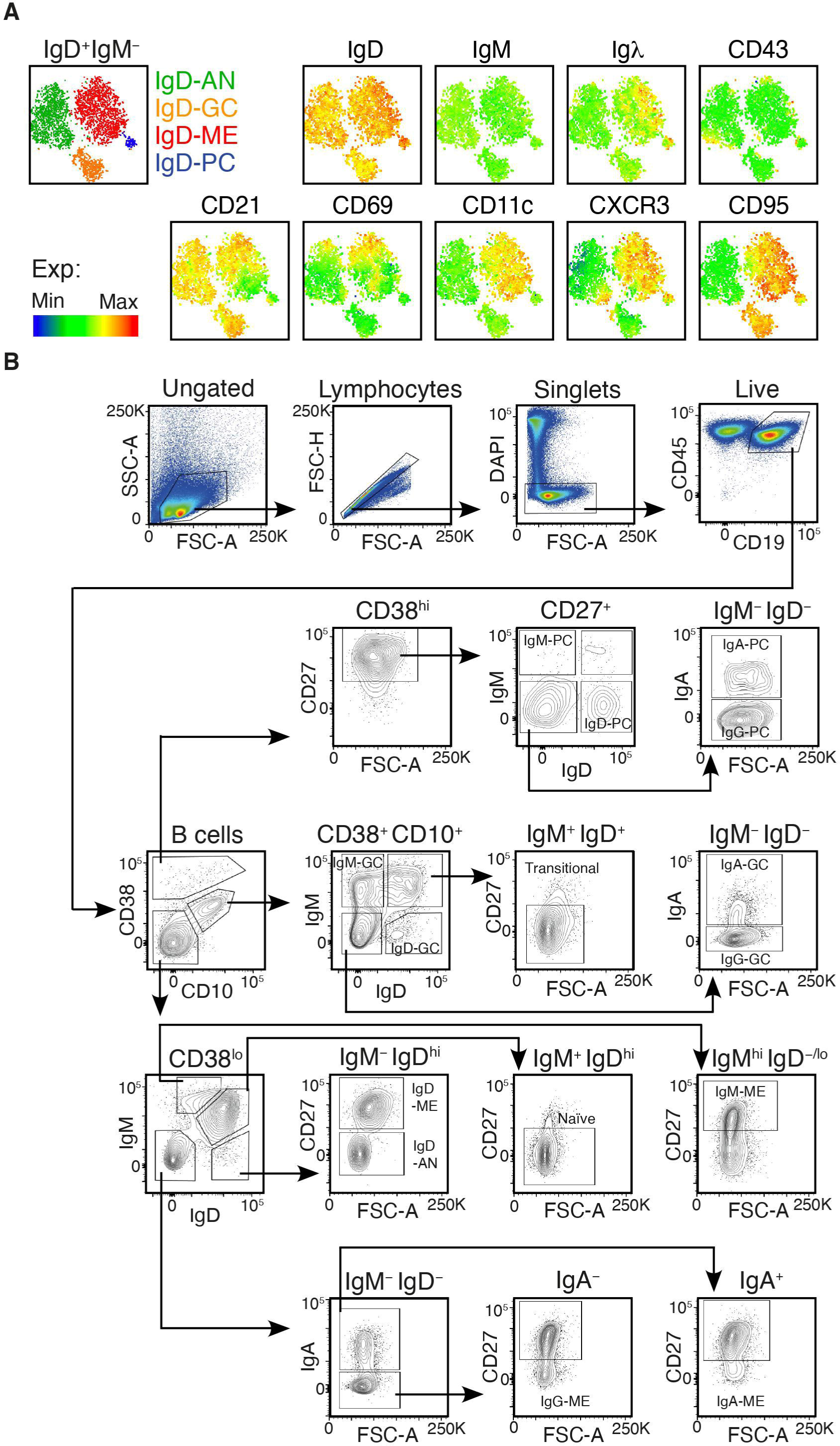
Phenotype and gating strategies to identify tonsillar B cell subsets. (**A**) Spectral flow cytometry tSNE plots of tonsillar IgD^+^IgM^−^ B cells showing clusters and relative expression of IgD, IgM, Igλ, CD43, CD21, CD69, CD11c, CXCR3 and CD95. Representative data from three independent experiments. (**B**) Gating strategy used to identify human IgM^−^IgD^+^ (IgD), IgM^+^IgD^−^ (IgM), IgM^−^IgD^−^IgA^−^ (IgG), and IgM^−^IgD^−^IgA^+^ (IgA) subsets from CD38^−^CD10^−^ ME B cells, CD38^+^CD10^+^ GC B cells or CD38^high^CD27^high^ PCs gated among total CD45^+^CD19^+^ B cells from human tonsils. Naive CD38^−^CD10^−^IgM^+^IgD^high^CD27^−^ B cells, AN CD38^−^CD10^−^IgM^−^IgD^high^ CD27^−^ B cells, and transitional IgM^+^IgD^+^CD38^+^ CD10^+^CD27^−^ B cells are also shown.

**Fig. S2.**
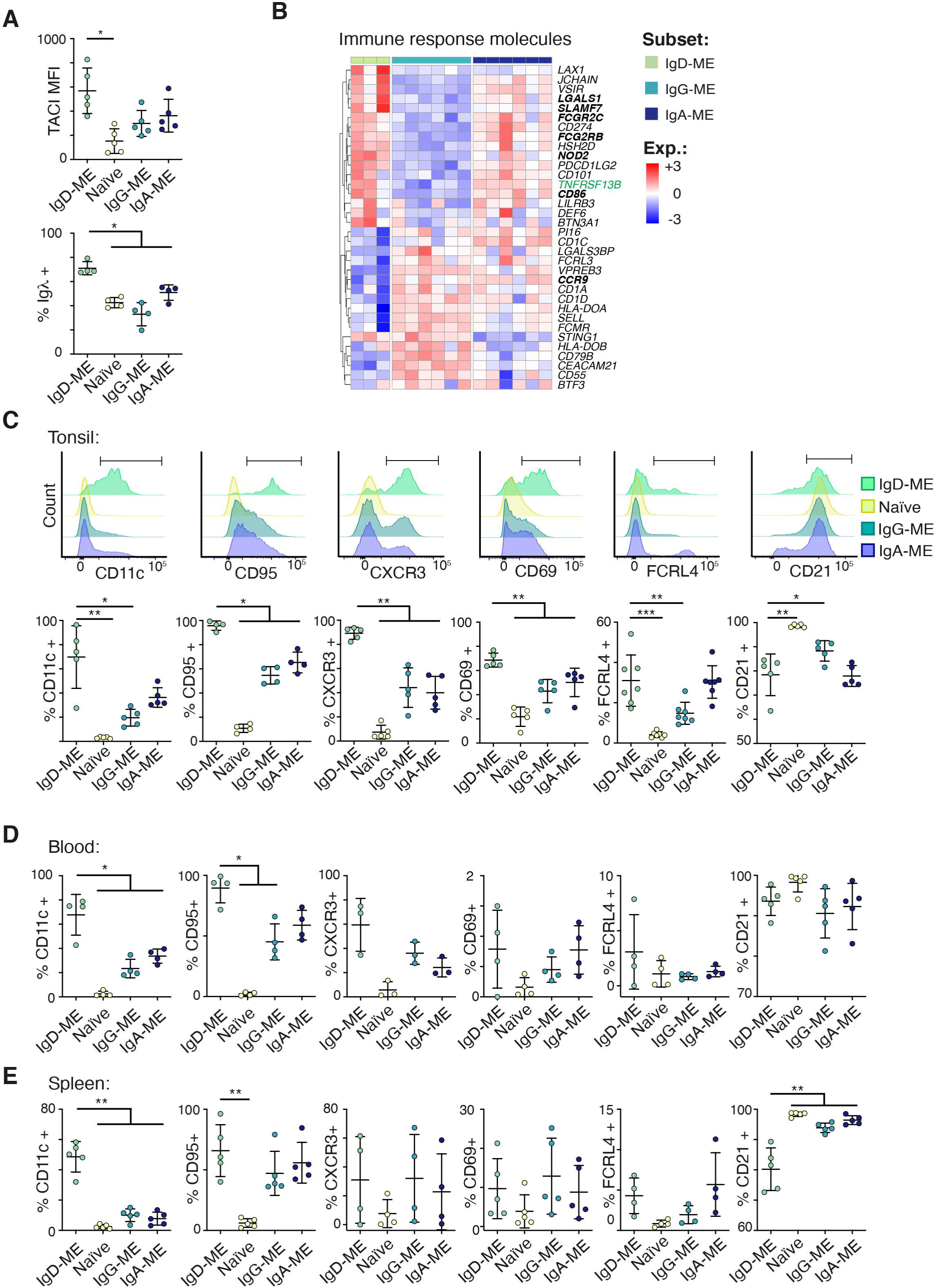
Tonsillar IgD-ME B cells exhibit unique tissue-specific phenotypic properties. (**A**) Flow cytometry analysis showing expression of TACI and surface Igλ across naïve, IgD-ME, IgG-ME or IgA-ME B cells from human tonsils (n = 4-5). (**B**) Heatmap showing differentially expressed genes (adj.P.Value < 0.05 and |log2FC| > 1) by IgD-ME vs. IgG-ME or IgA-ME B cells belonging to manually curated “Immune response molecules”. The color bar depicts normalized intensity values. Genes highlighted in bold are discussed in the text. (**C**) Representative flow cytometry profiles (top row) and summary graphs (bottom row) of CD11c, CD95, CXCR3, CD69, FCRL4 and CD21 expression on naïve, IgD-ME, IgG-ME or IgA-ME B cells from human tonsils (n = 3-6). (**D, E**) Summary graphs of surface markers detected as in (C) from circulating (D) or splenic (E) B cells. Error bars represent S.D. Differences were assessed with Kruskal-Wallis test followed by a post-hoc pairwise Mann-Whitney test. * p < 0.05, ** p < 0.01 and *** p < 0.001. Statistical significance was only calculated between IgD-ME B cells and other subsets. Comparisons lacking statistical reporting are not statistically significant.

**Fig. S3.**
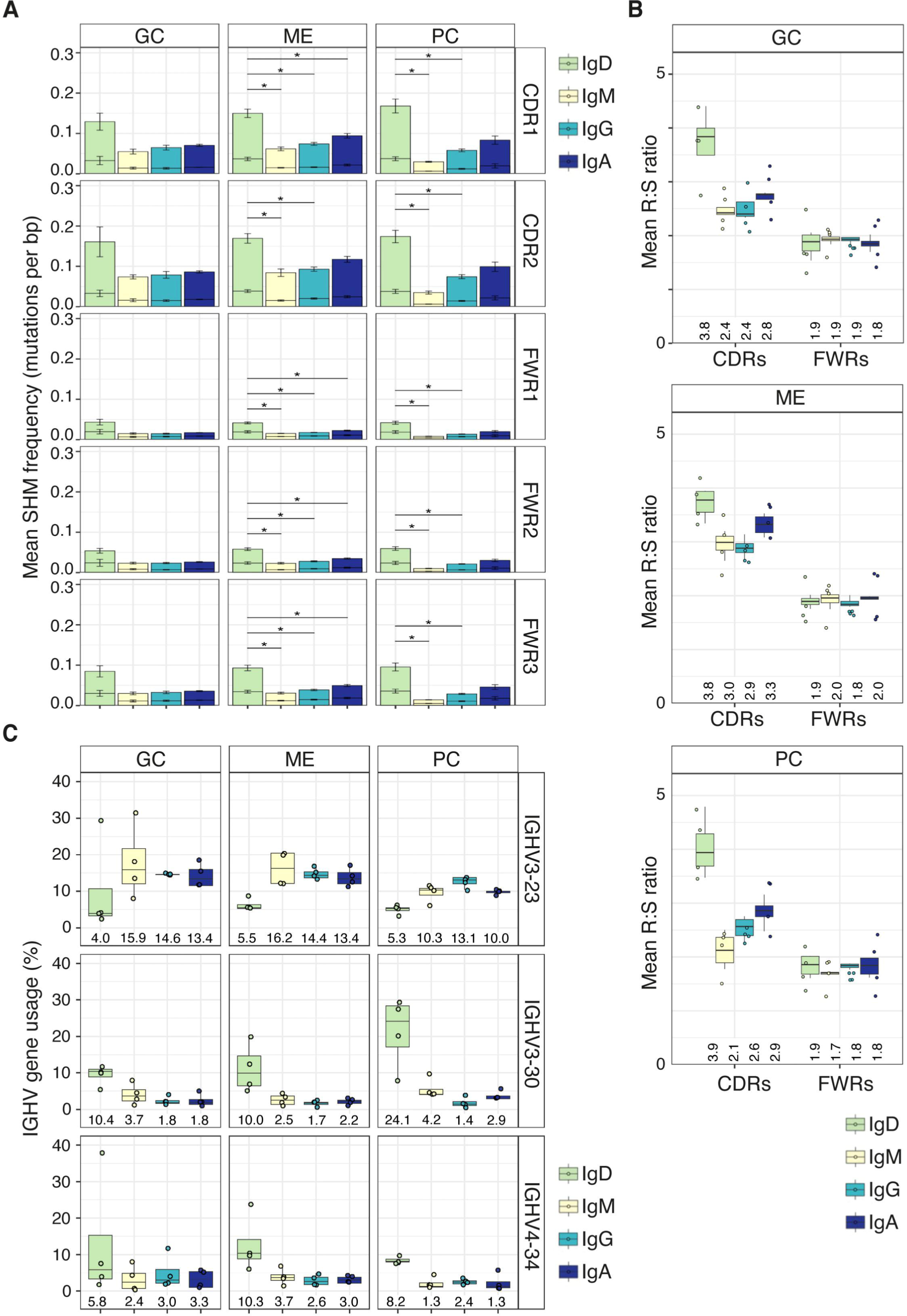
Tonsillar IgD-ME B cells and IgD-PCs display a mutation profile consistent with GC ontogeny. (**A**) Mean SHM frequency (mutations per bp) across CDRs and FWRs of IGHV genes from human tonsillar ME, GC, and PC subsets of all donors (n = 4). Within each bar, the top segment represents replacement mutation frequency, whereas the bottom segment represents silent mutation frequency. Error bars represent SEM. Differences in total mutational frequency were assessed with Kruskal-Wallis test followed by a post-hoc pairwise Mann-Whitney test. (**B**) Mean replacement-to-silent (R:S) mutation ratio across CDRs and FWRs of mutated antibodies expressed by tonsillar GC B cells, ME B cells or PC expressing only IgD, IgM, IgG or IgA (n=4). (**C**) Mean IGHV3-23, IGHV3-30 and IGHV4-34 gene usage by GC B cells, ME B cells and PC from human tonsils expressing surface IgM, IgD, IgG or IgA alone (n = 4). Numbers below bars in (B-C) represent median values for the four data points. Differences in (B-C) were assessed with Kruskal-Wallis test followed by a post-hoc pairwise Mann-Whitney test with p-value adjustment following the Benjamini-Hochberg method. * p-value < 0.05. Panels without significance bars did not report significant differences.

**Fig. S4.**
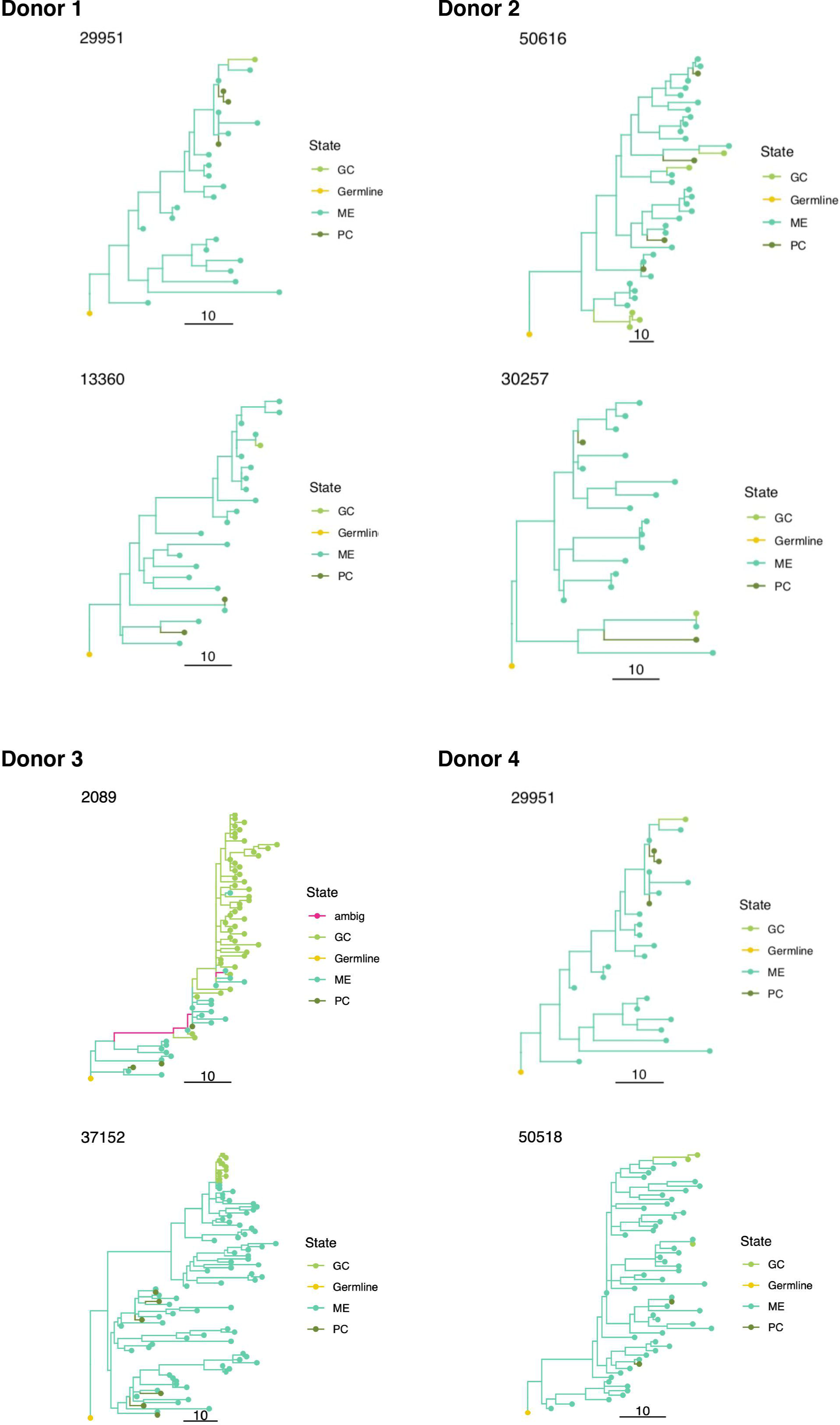
Tonsillar IgD-ME B cells participate in mutation-intensive and large lineage trees, and show more IgD-specific clonal relatedness to IgD-PC B cells than to IgD-GC B cells. Additional examples of lineage trees from the IgD gene repertoire of tonsillar IgD class-switched B cells from all study donors. The length of scale bars is equal to ten somatic hypermutations. GC; germinal centre, ME; memory, PC; plasma cell, ambig; ambiguity.

**Fig. S5.**
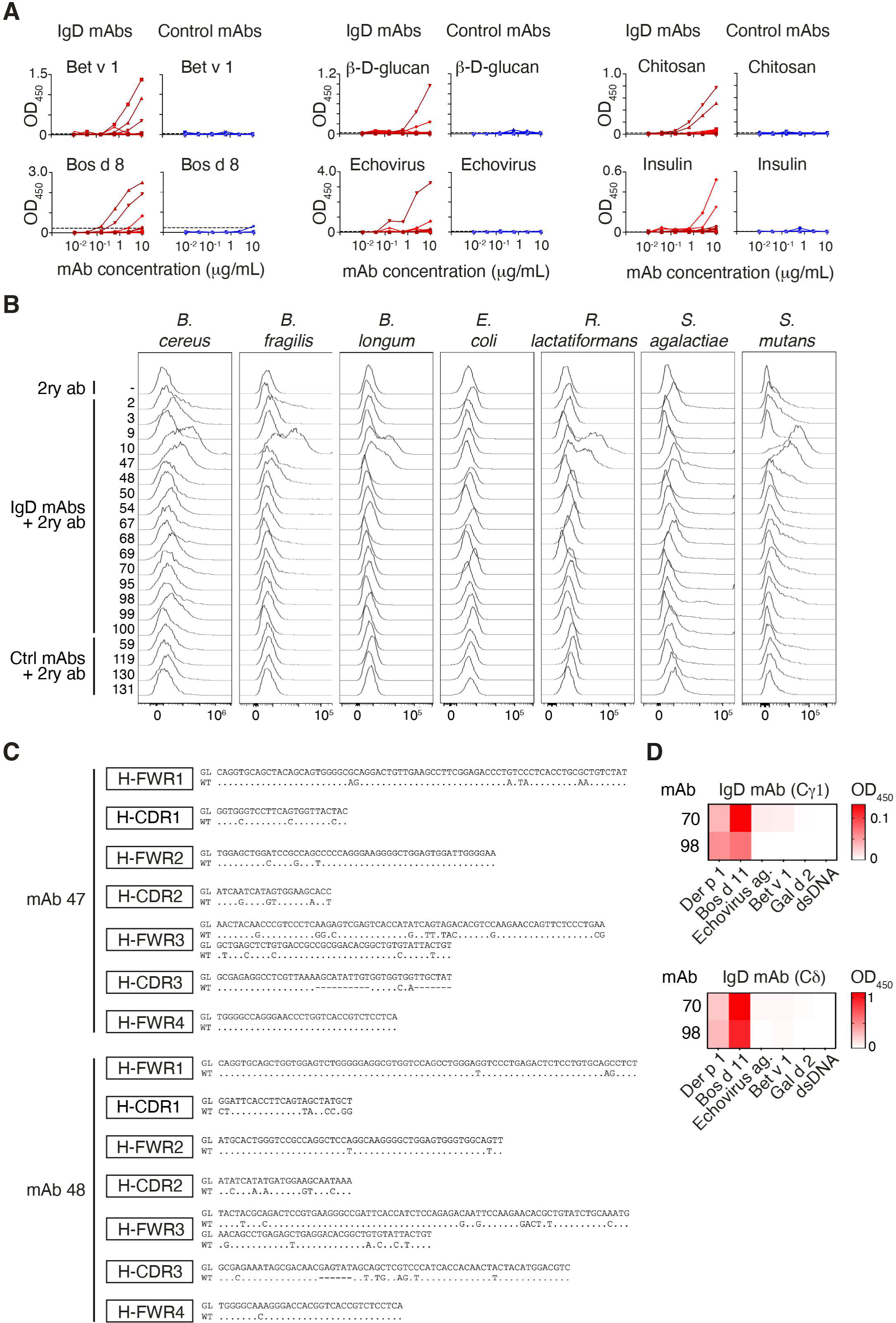
A subset of polyreactive IgD mAbs broadly reacts against common aerodigestive antigens, self-antigens and bacteria. (**A**) Binding curves of 23 IgD mAbs from tonsillar IgD-GC B cells (red) and 10 control mAbs from circulating B cells with known reactivity to SARS-CoV-2 (blue) to food, fungal, protist, viral, and polyreactivity-associated antigens measured by ELISA. Dashed lines indicate reactivity thresholds. (**B**) Flow cytometry histograms showing binding profiles of IgD mAbs, control (Ctrl) mAbs, or control secondary antibody (2ry ab) to isolated bacterial strains. (**C**) Variable IGHV4-34 and IGHV3-30 gene sequence from wild type (WT) mAbs 47 and 48, respectively, and their germline (GL) counterparts. Dashes represent missing GL nucleotides. (**D**) Heat maps showing ELISA binding intensity to selected antigens by 10 μg/mL mAbs 70 and 98 encompassing a Cγ1 (top) or Cδ (bottom) HC.

**Table S3.** Antibodies used for flow cytometry and cell sorting. ctlg#, catalogue number; RRID, research resource identifier; cFC, conventional flow cytometry; sFC, spectral flow cytometry; CS, cell sorting.

| **Antibody** | **Reference** | **Use** |
| --- | --- | --- |
| Alexa Fluor(R) 700 anti-human CD45 (clone: HI30) | BioLegend; ctlg# 304024; RRID: AB_493761 | cFC, CS |
| PE/Cyanine7 anti-human CD19 antibody (clone: HIB19) | BioLegend; ctlg# 302216; RRID: AB_14246 | cFC, CS |
| APC/Cyanine7 anti-human CD38 antibody (clone: HIT2) | BioLegend; ctlg# 303534; RRID: AB_2561605) | cFC, CS |
| APC anti-human CD10 antibody (clone: HI10a) | BioLegend; ctlg# 312210; RRID: AB_314921 | cFC, CS |
| PerCP/Cyanine5.5 anti-human CD27 antibody (clone: M-T271) | BioLegend; ctlg# 356407; RRID: AB_2561905 | cFC |
| Anti-Human CD27 Monoclonal Antibody, Phycoerythrin (PE) Conjugated (Clone: O323) | Thermo Fisher Scientific; ctlg# 12-0279-73; RRID: AB_465617 | CS |
| Brilliant Violet 605(TM) anti-human IgM antibody (clone: MHM-88) | BioLegend; ctlg# 314524; RRID: AB_2562374 | cFC, sFC, CS |
| PE-CF594 Mouse Anti-Human IgD (clone: IA6-2) | BD Biosciences; ctlg# 562540; RRID: AB_11153129 | cFC, sFC, CS |
| Anti-IgA-FITC, human antibody (clone: IS11-8E10) | Miltenyi Biotec; ctlg# 130-099-107; RRID: AB_2659715 | cFC, CS |
| PE anti-human CD1d antibody (clone: 51.1) | BioLegend; ctlg# 350306; RRID: AB_10641845 | cFC |
| PE anti-human CD11b antibody (clone: ICRF44) | BioLegend; ctlg# 301305; RRID: AB_314157 | cFC |
| PE Mouse Anti-Human CD11c (clone: B-ly6) | BD Biosciences; ctlg# 555392; RRID: AB_395793 | cFC |
| PE Mouse Anti-Human CD21 (clone: B-ly4) | BD Biosciences; ctlg# 555422; RRID: AB_395816 | cFC |
| anti-human CD62L PE-conjugated (clone: LT-TD180) | EuroBioscience; ctlg# H12201P | cFC |
| Anti-Human CD69 Monoclonal Antibody, Phycoerythrin (PE) Conjugated (Clone: FN50) | Thermo Fisher Scientific; ctlg# 12-0699-73; RRID: AB_465735 | cFC, sFC |
| PE anti-human CD71 antibody (clone: CY1G4) | BioLegend; ctlg# 334105; RRID: AB_2271603 | cFC |
| Mouse Anti-CD73 Monoclonal Antibody, Phycoerythrin Conjugated (Clone: AD2) | BD Biosciences; ctlg# 550257; RRID: AB_393561 | cFC |
| Mouse Anti-CD80 Monoclonal Antibody, Phycoerythrin Conjugated (Clone: L307.4) | BD Biosciences; ctlg# 557227; RRID: AB_396606 | cFC |
| PE anti-human CD95 (Fas) antibody (clone: DX2) | BioLegend; ctlg# 305607; RRID: AB_314545 | cFC |
| PE anti-human CD183 (CXCR3) antibody (clone: G025H7) | BioLegend; ctlg# 353705; RRID: AB_10959652 | cFC |
| PE anti-human CD307d (FcRL4) antibody (clone: 413D12) | BioLegend; ctlg# 340203; RRID: AB_1575103 | cFC |
| PE anti-human CD307e (FcRL5) antibody (clone: 509f6) | BioLegend; ctlg# 340304; RRID: AB_2104588 | cFC |
| PE anti-human Ig light chain lambda antibody (clone: MHL-38) | BioLegend; ctlg# 316607; RRID: AB_493626 | cFC |
| Human TACI/TNFRSF13B Phycoerythrin Mab (clone: 165604) | R&D Systems; ctlg# FAB1741P; RRID: AB_2203409 | cFC |
| Brilliant Violet 421(TM) anti-human CD10 antibody (clone: HI10a) | BioLegend; ctlg# 312217; RRID: AB_10899409 | sFC |
| Pacific Blue(TM) anti-human CD19 antibody (clone: HIB19) | BioLegend; ctlg# 302224; RRID: AB_493653 | sFC |
| BV480 Mouse Anti-Human Ig, $\lambda$ Light Chain (clone: 1-155-2) | BD Biosciences; ctlg# 751013; RRID: AB_2875061 | sFC |
| BV650 Mouse Anti-Human CD11c antibody (clone: B-ly6) | BD Biosciences; ctlg# 563403; RRID: AB_2732048 | sFC |
| BV711 Mouse Anti-Human CD43 (clone: 1G10) | BD Biosciences; ctlg# 743614; RRID: AB_2741624 | sFC |
| BV786 Mouse Anti-Human CD21 (clone: B-ly4) | BD Biosciences; ctlg# 740969; RRID: AB_2740594 | sFC |
| CD45 Monoclonal Antibody (HI30), Alexa Fluor 532 (clone: HI30) | Thermo Fisher Scientific; ctlg# 58-0459-41; RRID: AB_11218084 | sFC |
| IgA1-PerCP-Cyanine5.5 (clone: SAA1) | CYTOGNOS; ctlg#_CYT-IGA1C | sFC |
| IgA2-PerCP-Cyanine5.5 (clone: SAA2) | CYTOGNOS; ctlg# CYT-IGA2C2 | sFC |
| PE-Cy<sup>TM</sup>5 Mouse Anti-Human CD183 (clone: 1C6/CXCR3) | BD Biosciences; ctlg# 561731; RRID: AB_10892799 | sFC |
| PE/Cyanine7 anti-human CD95 (Fas) antibody (clone: DX2) | BioLegend; ctlg# 305621; RRID: AB_2100370 | sFC |
| APC mouse anti-human CD27 (clone: M-T271) | BD Biosciences; ctlg# 561786, RRID: AB_10896653 | sFC |
| Anti-human CD38 APC Fire 810 (clone: HB-7) | BioLegend; ctlg# 356643; RRID: AB_2860936 | sFC |

**Table S4.** Antibodies used for tissue immunofluorescence assays. ctlg#, catalogue number; RRID, research resource identifier; 1ry, primary; 2ry, secondary.

| **Antibody** | **Reference** | **Use** |
| --- | --- | --- |
| FLEX Polyclonal Rabbit Anti-Human IgD Ready-to-Use (polyclonal) | Dako; ctlg# IR517 | 1ry |
| Mouse Anti-Human IgD-UNLB antibody (clone: IADB6) | SouthernBiotech; ctlg# 9030-01; RRID: AB_2796590 | 1ry |
| Goat anti-Human IgM Fc Antibody (polyclonal) | Thermo Fisher Scientific; ctlg# H15000; RRID: AB_2536556 | 1ry |
| Rabbit anti- Pan-Cytokeratin (H-240) antibody (polyclonal) | Santa Cruz Biotechnology; ctlg# sc-15367; RRID: AB_2134438 | 1ry |
| Rat Ki-67 Monoclonal Antibody (SolA15), eBioscience™ (clone: SolA15) | Thermo Fisher Scientific; ctlg #14-5698-82, RRID: AB_10854564 | 1ry |
| IgA1 Antibody (clone: H-11) | Santa Cruz Biotechnology; ctlg# sc-271913, RRID:AB_10609520 | 1ry |
| Anti-IgA2 antibody (polyclonal) | Abcam; ctlg# ab88250, RRID:AB_2041786 | 1ry |
| Alexa Fluor® 488 AffiniPure F(ab')<sub>2</sub> Fragment Donkey Anti-Rabbit IgG (H+L) (polyclonal) | Jackson ImmunoResearch Labs; ctlg# 711-546-152; RRID: AB_2340619 | 2ry |
| Alexa Fluor® 488 AffiniPure F(ab')<sub>2</sub> Fragment Donkey Anti-Rat IgG (H+L) (polyclonal) | Jackson ImmunoResearch Labs; ctlg# 712-546-153; RRID: AB_2340686 | 2ry |
| Cy™3 AffiniPure F(ab')<sub>2</sub> Fragment Donkey Anti-Goat IgG (H+L) (polyclonal) | Jackson ImmunoResearch Labs; ctlg# 705-166-147; RRID: AB_2340413 | 2ry |
| Cy™3 -AffiniPure F(ab')<sub>2</sub> Fragment Donkey Anti-Rabbit IgG (H+L) antibody (polyclonal) | Jackson ImmunoResearch Labs; ctlg# 711-166-152; RRID: AB_2313568 | 2ry |
| Alexa Fluor 647-AffiniPure Donkey Anti-Rabbit IgG (H+L) | Jackson ImmunoResearch Labs; ctlg# 711-605-152, RRID:AB_2492288 | 2ry |
| Cy<sup>TM</sup>5 AffiniPure Donkey Anti-Mouse IgG (H+L) (polyclonal) | Jackson ImmunoResearch Labs;<br>ctlg# 715-175-151; RRID:<br>AB_2340820 | 2ry |
| Alexa Fluor® 647 AffiniPure F(ab')<sub>2</sub> Fragment Donkey Anti-Goat IgG (H+L) (polyclonal) | Jackson ImmunoResearch Labs;<br>ctlg# 705-606-147; RRID:<br>AB_2340438 | 2ry |
| Alexa Fluor 488-AffiniPure Donkey Anti-Mouse IgG (H+L) | Jackson ImmunoResearch Labs<br>ctlg# 715-545-150,<br>RRID:AB_2340846 | 2ry |

**Table S5.**
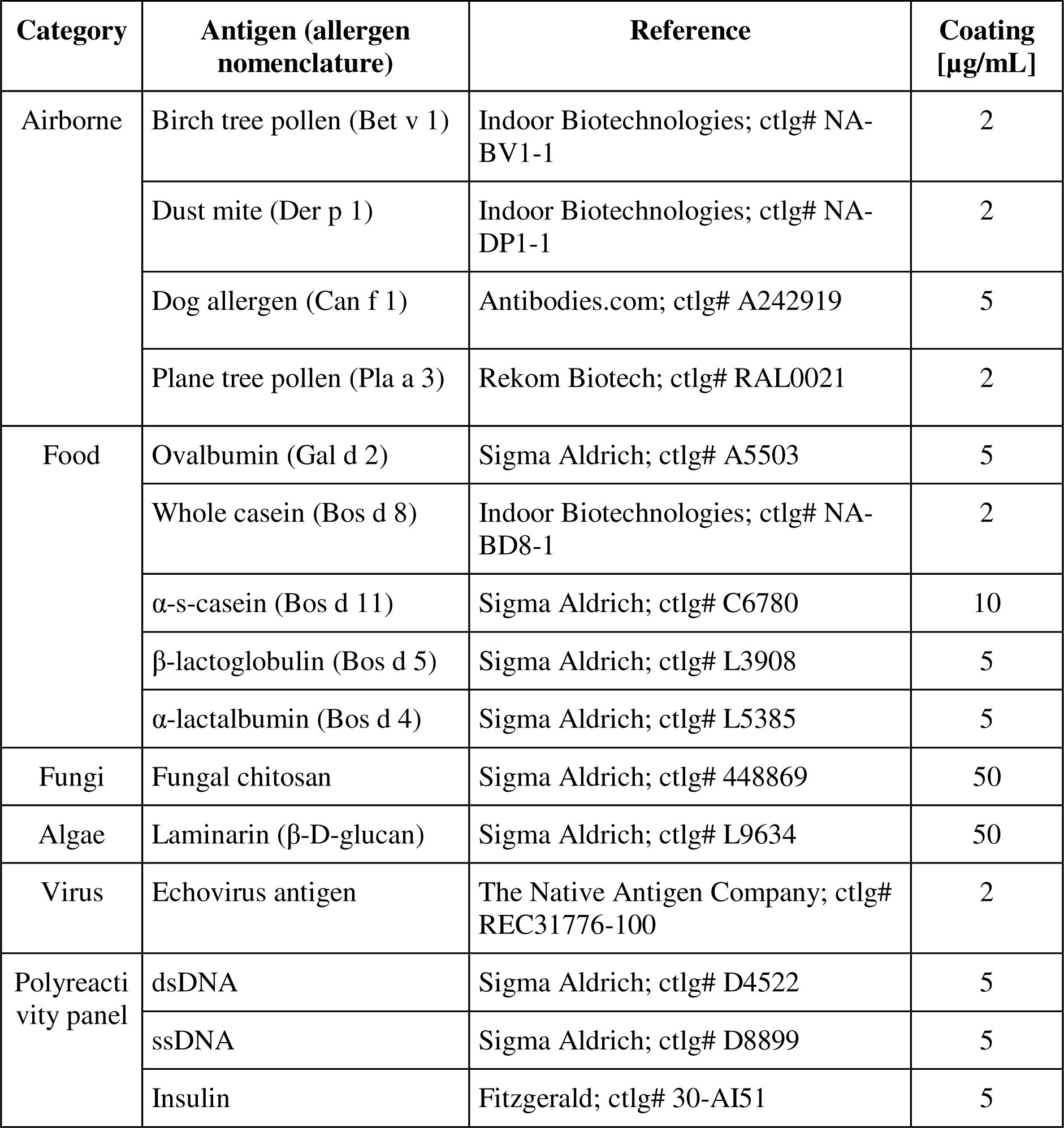
Antigens used in ELISAs measuring antibody reactivity. ctlg#, Catalogue number.

| Category | Antigen (allergen nomenclature) | Reference | Coating [µg/mL] |
| --- | --- | --- | --- |
| Airborne | Birch tree pollen (Bet v 1) | Indoor Biotechnologies; ctlg# NA-BV1-1 | 2 |
|  | Dust mite (Der p 1) | Indoor Biotechnologies; ctlg# NA-DP1-1 | 2 |
|  | Dog allergen (Can f 1) | Antibodies.com; ctlg# A242919 | 5 |
|  | Plane tree pollen (Pla a 3) | Rekom Biotech; ctlg# RAL0021 | 2 |
| Food | Ovalbumin (Gal d 2) | Sigma Aldrich; ctlg# A5503 | 5 |
|  | Whole casein (Bos d 8) | Indoor Biotechnologies; ctlg# NA-BD8-1 | 2 |
|  | α-s-casein (Bos d 11) | Sigma Aldrich; ctlg# C6780 | 10 |
|  | β-lactoglobulin (Bos d 5) | Sigma Aldrich; ctlg# L3908 | 5 |
|  | α-lactalbumin (Bos d 4) | Sigma Aldrich; ctlg# L5385 | 5 |
| Fungi | Fungal chitosan | Sigma Aldrich; ctlg# 448869 | 50 |
| Algae | Laminarin (β-D-glucan) | Sigma Aldrich; ctlg# L9634 | 50 |
| Virus | Echovirus antigen | The Native Antigen Company; ctlg# REC31776-100 | 2 |
| Polyreactivity panel | dsDNA | Sigma Aldrich; ctlg# D4522 | 5 |
|  | ssDNA | Sigma Aldrich; ctlg# D8899 | 5 |
|  | Insulin | Fitzgerald; ctlg# 30-AI51 | 5 |

**Table S6.**
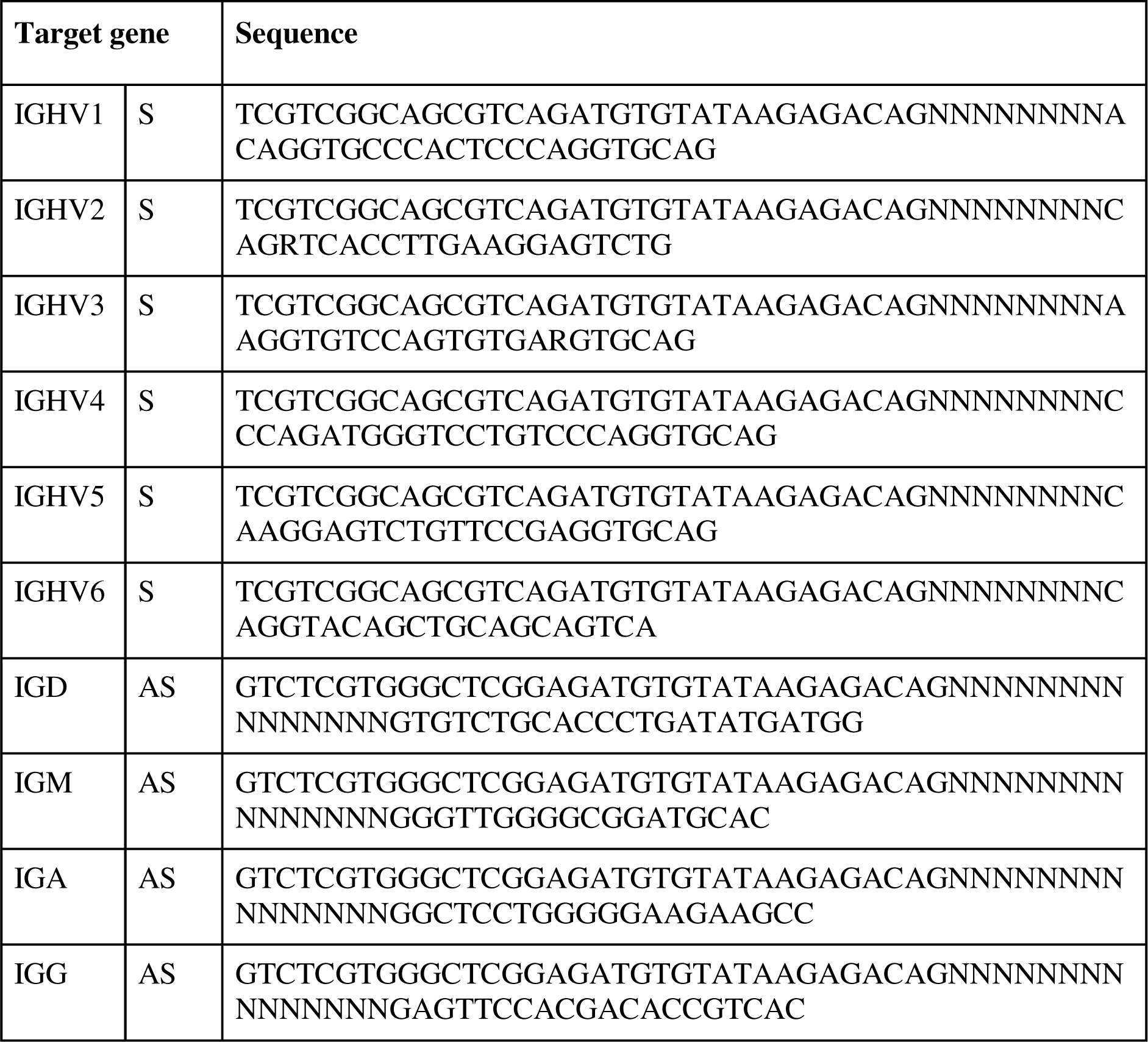
Primers used for Ig gene repertoire sequencing. S; sense, AS; antisense.

